# Estradiol regulates local synthesis of synaptic proteome via sex-specific mechanisms

**DOI:** 10.1101/2023.12.17.571898

**Authors:** Pooja Raval, Hannah Rae, Rodrigo R. R. Duarte, Iain A. Watson, Katherine J. Sellers, Kathryn M. C. Pachon, Laura Sichlinger, Timothy R. Powell, Marina V. Yasvoina, Jayanta Mukherjee, Stephen J. Moss, Nicholas J. Brandon, Deepak P. Srivastava

**Affiliations:** Maurice Wohl Clinical Neuroscience Institute, Department of Basic and Clinical Neuroscience, Institute of Psychiatry, Psychology, & Neuroscience, King’s College London, London, UK; MRC Centre for Neurodevelopmental Disorders, King’s College London, London, UK; Social, Genetic & Developmental Psychiatry Centre, Institute of Psychiatry, Psychology & Neuroscience, King’s College London, London, UK; Department of Neuroimaging, Institute of Psychiatry, Psychology and Neuroscience, King’s College London, London, UK; AstraZeneca-Tufts Laboratory for Basic and Translational Neuroscience, Tufts University Medical School, Boston, MA, USA; Discovery, Neuroscience, BioPharmaceuticals R&D, AstraZeneca, Boston, MA, USA

## Abstract

Estrogens, specifically 17β-estradiol (estradiol), can modulate synaptic function by regulating the expression and localisation of synaptic proteins. However, the mechanisms underlying estradiol’s regulation of synaptic protein expression, and whether if they occur in a sex specific manner, is not well understood. In this study, using sex-specific hippocampal slice cultures and mixed-sex primary hippocampal neurons, we investigated whether local protein synthesis is required for estradiol-induced synaptic protein expression. Estradiol rapidly increased the rate of protein synthesis and the number of actively translating ribosomes along dendrites and near synapses in both male and female hippocampal neurons. Importantly, these effects occurred independently of gene transcription. Moreover, estradiol also increased the abundance of nascent proteins localised to synapses, independently of gene transcription. Specifically, estradiol increased the synaptic expression of GluN2B- containing *N*-methyl-D-aspartate receptors and PSD-95 in male and female hippocampus. Mechanistically, mTOR signalling was required for estradiol-induced increases in overall local protein synthesis only in male but not female hippocampus. Consistent with this, mTOR signalling mediated estradiol increases in GluN2B in male, but not female, hippocampus. Conversely, mTOR inhibition, blocked estradiol-induced increased PSD-95 expression in both male and female hippocampus. Collectively, these data suggest that the rapid modulation of local protein synthesis by estradiol is required for changes in the synaptic proteome in male and female hippocampus, and that the requirement of the mTOR signalling pathway in these effects occur in both a sex-specific and protein-dependent manner, with this signalling pathway have a greater role in male compared to female hippocampus.

## Introduction

Estrogens, in particular the biologically active form 17β-estradiol (estradiol; E2), have widely been reported to have an enhancing effect on cognition in rodents (Taxier *et al*., 2020). A single administration of estradiol is sufficient to improve performance on a range of cognitive tasks in both male and female rodents (Fortress *et al*., 2013; Phan *et al*., 2015; Jacome *et al*., 2016). These effects occur in parallel to estradiol-induced dendritic spine formation (Phan *et al*., 2015; Tuscher *et al*., 2016; Luine *et al*., 2018), and increased expression of excitatory synaptic proteins. For example, estradiol increases the synaptic expression of GluN2B-containing *N*-methyl-D- aspartate receptors (NMDARs), PSD-95 and GluA1-containing α-amino-3-hydroxy-5- methyl-4-isoxazoleproprionic acid receptors (AMPARs) (Jelks *et al*., 2007; Liu *et al*., 2008; Smith *et al*., 2009; Smith *et al*., 2016; Avila *et al*., 2017; Sheppard *et al*., 2023). These changes in dendritic spine density and synaptic proteome are thought to underlie estradiol’s ability to modulate synaptic transmission and plasticity, and thus, its effects on cognition (Srivastava *et al*., 2013; Vedder *et al*., 2013; Luine *et al*., 2018; Sheppard *et al*., 2019). Indeed, estradiol rapidly potentiates excitatory synaptic transmission the hippocampus (Wong & Moss, 1992; Oberlander & Woolley, 2016; Jain & Woolley, 2023), and enhances *N*-methyl-D-aspartate receptor (NMDAR)- dependent long-term potentiation (Smith & McMahon, 2006; Liu *et al*., 2008; Kramar *et al*., 2013; Vedder *et al*., 2013). Interestingly, while these effects occur in both male and female hippocampus, the mechanisms underlying estradiol’s effects on synaptic transmission differ in a sex-specific manner (Oberlander & Woolley, 2016; Jain & Woolley, 2023). However, the mechanism underlying estradiol’s effect on the synaptic proteome, and whether they also occur in a sex-specific manner is not well understood.

Mechanistically, the effects of estradiol on the synaptic proteome, dendritic spines and synaptic function are thought to be mediated via a ‘non-classical’ membrane-initiated mechanism (Srivastava *et al*., 2013; Taxier *et al*., 2020). Estradiol-mediated membrane-initiated signalling is characterised by activation of signalling kinase pathways, such as the extracellular signal-regulated kinase (ERK) and mammalian target of Rapamycin (mTOR) pathways, as well as the rearrangement of the actin cytoskeleton, and moreover, is thought to be independent of gene transcription (Sellers *et al*., 2015a; Taxier *et al*., 2020). Consistent with this, activation of kinases pathways, as well as effects on dendritic spines, synaptic proteome and function have typically been described to occur within minutes of estradiol treatment (Woolley, 2007; Phan *et al*., 2015; Sellers *et al*., 2015a; Luine *et al*., 2018; Taxier *et al*., 2020; Sheppard *et al*., 2023). However, the effects of estradiol on the synaptic proteome, dendritic spines and synaptic function have also been described to last for hours or days (Woolley & McEwen, 1994; Smith & McMahon, 2006; Liu *et al*., 2008; Tuscher *et al*., 2016). As such, it is currently not fully understood how estradiol treatment can result in long-lasting changes in the synaptic proteome and synaptic function.

The changes in the synaptic proteome that drive synaptic plasticity can be achieved by the local synthesis of specific proteins along dendrites and at synapses. Consistent with, the machinery required for local synthesis of protein as well as mRNA encoding for synaptic proteins have been found to be located along dendrites and near synapses (Shrestha & Klann, 2022; Bourke *et al*., 2023). Moreover, neurons have been shown to increase the expression of synaptic proteins via a local protein synthesis mechanism, in response to different stimuli, including synaptic (Huber *et al*., 2000; Schanzenbacher *et al*., 2016; Hafner *et al*., 2019) and neuromodulatory signals including brain-derived neurotrophic factor (BDNF) (Hafner *et al*., 2019; Jung *et al*., 2020) and dopamine (Smith *et al*., 2005; David *et al*., 2020). Therefore, local protein synthesis provides neurons with a mechanism to rapidly refine the synaptic proteome independently of gene transcription. A signalling pathway that has received particular attention in mediating local protein synthesis is the mTOR pathway (Costa-Mattioli *et al*., 2009; Hoeffer & Klann, 2010). mTOR has well described roles in memory enhancement (Bekinschtein *et al*., 2007), and has also been shown to be critical for estradiol-dependent enhancement in the object recognition memory task (Fortress *et al*., 2013). Furthermore, estradiol rapidly activates mTOR *in vivo* (Fortress *et al*., 2013) and *in vitro* (Sarkar *et al*., 2010; Briz & Baudry, 2014; Sellers *et al*., 2015b) and is required for the maintenance of estradiol-induced increased spine density (Tuscher *et al*., 2016), but not formation (Sellers *et al*., 2015b). This has led to the suggestion that estradiol may influence local protein synthesis. In line with this, estradiol regulates the activity of mTOR downstream targets, eukaryotic initiation factor 4E (4EBP1), p70 ribosomal S6 kinase (p70S6K) and ribosomal protein S6 (RPS6) (Akama & McEwen, 2003; Sarkar *et al*., 2010; Fortress *et al*., 2013), which regulate local protein synthesis in neurons (Hoeffer & Klann, 2010; Rangaraju *et al*., 2017). Consistent with these observations, estradiol has been shown to rapidly increase dendritic mRNA translation through the use of a fluorescent reporter based on the fusion of GFP with the 3’ untranslated region (UTR) of calcium/calmodulin-dependent kinase IIα (Sarkar *et al*., 2010). Importantly, inhibition of protein synthesis, but not gene transcription, has been shown to be required for estradiol-facilitation of cognition (Sheppard *et al*., 2021). However, it is currently unclear whether estradiol modulation of synaptic proteome and dendritic spine density could occur via a local protein synthesis mechanism in male and female hippocampus, and if so, whether this requires signalling via the mTOR pathway.

In this study, we directly tested if estradiol could rapidly increase protein synthesis independently of gene transcription using a combination of acute hippocampal slice preparations from male and female mice, as well as mixed sex primary neuronal cultures. We further explored whether the rapid changes in dendritic spine density and synaptic proteome induced by estradiol were dependent on a local protein synthesis mechanism. Additionally, we investigated the role of the mTOR pathway in regulating local protein translation and changes in the synaptic proteome in male and female hippocampus. Data from our study suggests that the rapid modulation of local protein synthesis by estradiol is required for changes in the synaptic proteome in male and female hippocampus, but that the underlying signalling pathways that mediate these effects occur in a sex-specific manner.

## Results

### Estradiol increases local protein synthesis in the male and OVX female hippocampus

Estradiol has previously been demonstrated to rapidly engage protein translation machinery (Akama & McEwen, 2003; Sarkar *et al*., 2010; Fortress *et al*., 2013). However, whether estradiol regulates protein synthesis in a rapid time frame, either in male or female hippocampus, has not been directly tested. Therefore, we first assessed the rate of global protein synthesis following treatment with estradiol using the surface sensing of translation (SUnSET) assay (Schmidt *et al*., 2009) (**Supplementary Figure 1A-C**). Using mixed sex primary neuronal culture, we treated neurons with 10 nM estradiol for 15, 30, 60 or 120 minutes; the concentration of 10 nM has previously been shown to have significant effects on dendritic spine density and activate multiple kinase pathways (Erli *et al*., 2020). Puromycin (10 µg/mL) was added for the last 10 minutes of treatment (**Supplementary Figure 2A+B**). Estradiol increased the rate of protein synthesis which was significant after 120 minutes treatment (**Supplementary Figure 2B**). This time-point was used in subsequent experiments.

We next generated acute hippocampal slices from 10-12 week old male and ovariectomized (OVX) female mice, and treated them with estradiol (10 nM) for 2 hours; puromycin (5 µg/mL) was added for the final 30 minutes of the treatment (**Figure 1A**). Up to 4 hippocampal slices were cut from each brain. Slices were divided and used in two experimental designs: (1) vehicle (DMSO) vs. estradiol (E2); or (2) vehicle (DMSO) vs. estradiol (E2) vs. vehicle (DMSO) + inhibitor vs. estradiol (E2) + inhibitor. This experimental design allowed for a direct comparison of treatment to control conditions within the same hippocampal preparation (**Figure 1B**). Consistent with what we observed in primary cultures, estradiol treatment resulted in a significant increase in global protein synthesis in both male and OVX female hippocampal slices (**Figure 1C-F**).

**Figure 1.**
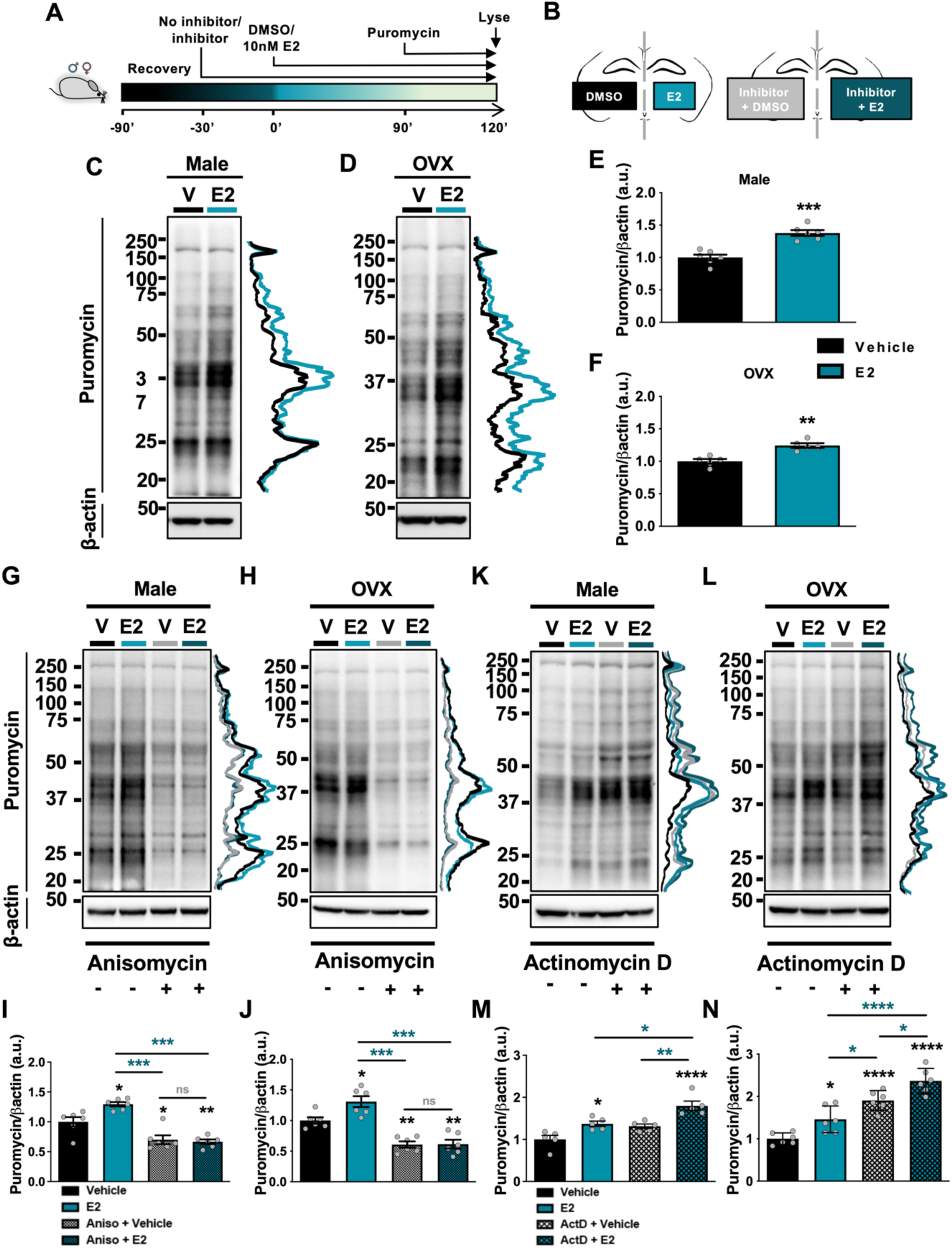
Estradiol increases protein synthesis independently of gene transcription in the male and OVX female hippocampus. **A** Timeline of pharmacological treatments **B** Diagram of the experimental set up: acute hippocampal slices were prepared from 10-12 week old male and ovariectomised (OVX) mice and treated with vehicle (DMSO; V) or estradiol (E2; 10nM), with or without protein translation inhibitor, anisomycin (Aniso, 40µM) or RNA polymerase II inhibitor, actinomycin D (ActD; 25µM) within the same animal. **C+D, G+H, K+L** Representative western blots of male **(C, G, K)** and OVX female **(D, H, L)** hippocampal slices pre-treated for 30 minutes with no inhibitor, anisomycin or actinomycin D, followed by estradiol (2 hours) or vehicle control (2 hours) and puromycin (SUnSET assay, 5µg/mL, last 30 minutes) treatment. Slices were processed for western blotting, immunoblotted for puromycin and normalised to housekeeper, β-actin. **E+F, I+J, M+N** Quantification of C+D, G+H, K+L. Estradiol acutely increased the rate of translation in male (**E**) and OVX female (**F**) hippocampus; unpaired student’s t-test, Welch’s correction; n=5/6. Anisomycin inhibited estradiol-mediated increases in the rate of protein synthesis in both male (**G**) and OVX female (**H**) hippocampus; one-way ANOVA, Bonferroni corrected; n=6. Whereas, estradiol continued to increase protein synthesis independently of actD inhibition of gene transcription in males (**M**) and OVX female (**N**) hippocampus; one-way ANOVA, Bonferroni corrected; n=6. Line-scan next to each western blot is a visual representation of intensity changes within puromycin bands. Error bars represent mean ± SEM; * p=<0.05, ** p = <0.01, *** p = <0.001, *** p = <0.001, ns = not significant.

Next, we determined whether estradiol-induced increase in global protein synthesis was a consequence of local protein synthesis. To this end, hippocampal slices were pre-treated with either the protein translation inhibitor, anisomycin or the RNA polymerase (transcriptional) inhibitor, actinomycin D (**Figure 1A**). Anisomycin blocked estradiol-mediated protein synthesis in both male and OVX female hippocampal slices (p=<0.0001; **Figure 1G-J**). Conversely, estradiol was still able to increase the rate of protein synthesis in the presence of actinomycin D in both male and OVX female hippocampal slices (p=<0.0001; **Figure 1K-N**). This indicates that estradiol was increasing the rate of hippocampal protein synthesis in a gene transcription independent manner in both sexes.

### Estradiol-induced protein translation localises to dendrites and dendritic spines

As estradiol treatment increased protein synthesis in a gene transcription-independent manner, we reasoned that it may be engaging translation machinery located along dendrites and within dendritic spines (Steward & Schuman, 2001). To test this, we first used primary hippocampal neurons and assessed where protein translation was occurring in neurons using SUnSET. Neurons were treated with estradiol (10 nM) or a vehicle for 2 hours; puromycin (10 ug/mL) was added for the last 10 minutes of treatment. Cell lysates were then fractionated to extra-nuclear and crude synaptic fractions. Estradiol treated neurons exhibited an increase in puromycylated polypeptide chains (proteins^PURO^) in the extra-nuclear fraction (p=0.0221; **Figure 2A+B**), but also in crude synaptic fractions (p=0.0009; **Figure 2A- D**), suggesting an increase in protein translation near synapses.

**Figure 2.**
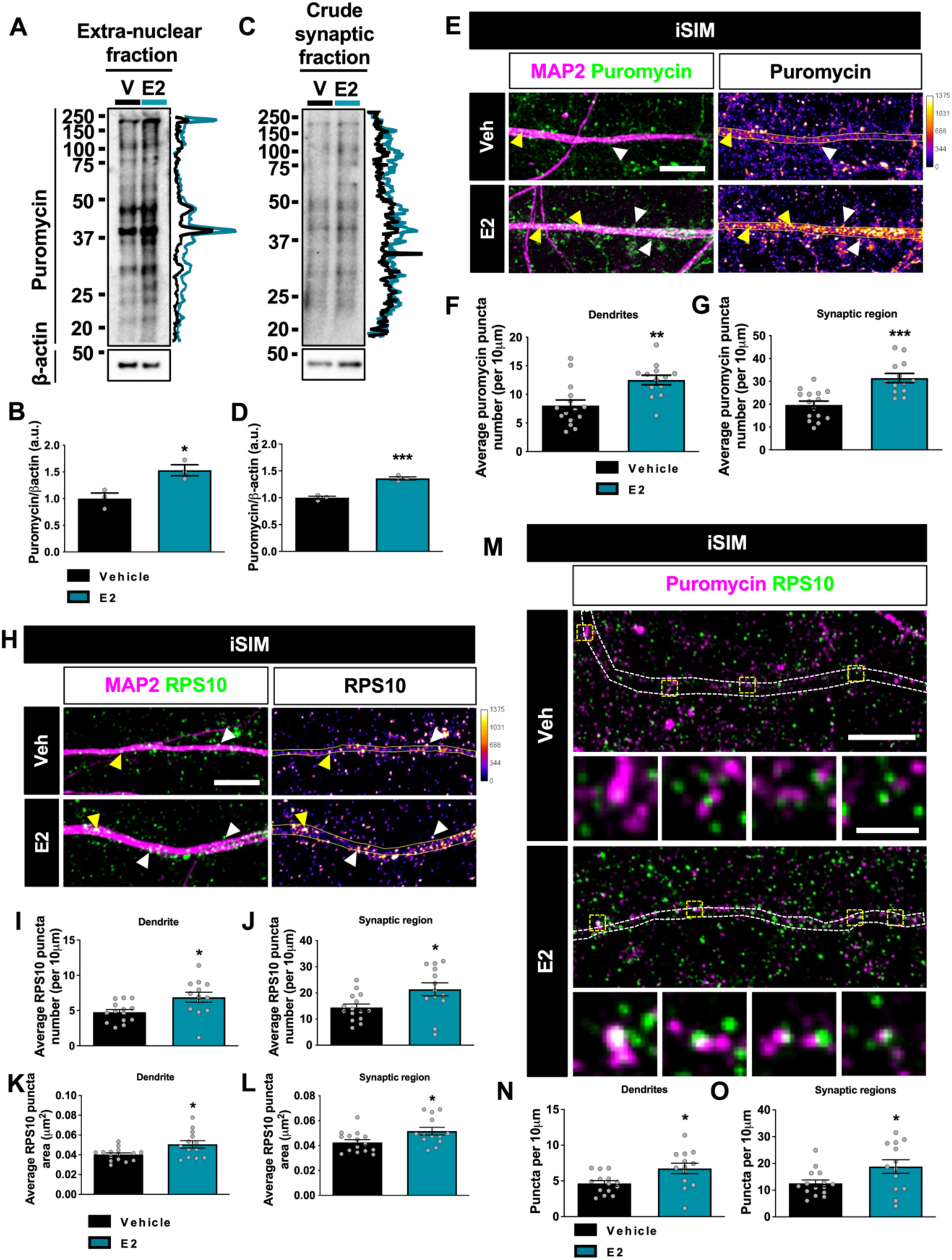
Estradiol increases protein synthesis near synapses. **A+C** Representative western blots of DIV 27 primary hippocampal neurons treated with estradiol (E2; 10nM, 2 hours) or vehicle control (DMSO; V; 2 hours) and puromycin (SUnSET assay, 10 µg/mL, last 10 minutes) fractionated upon lysing to separate out the extra-nuclear **(A)** and crude synaptosomal **(C)** fractions. Fractions were processed for western blotting, and immunoblotted for puromycin and normalised to housekeeper, β-actin. **C+D** Quantification of A+B. The rate of global protein synthesis was increased in estradiol-treated neurons in both extra-nuclear (**B**) and crude synaptosomal (**D**) fractions; n=3. **E, H, M** Representative super-resolution (iSIM) images of DIV 20 primary hippocampal neurons treated with estradiol or vehicle (2 hours) and puromycin (final 10 minutes). Neurons were immunostained for MAP2 (neuronal marker (magenta)) and puromycin (green; SUnSET-ICC) **(E)**; MAP2 (magenta) and component of 40S ribosomal subunit, ribosomal protein S10 (green; RPS10) **(H)**; puromycin (magenta) and RPS10 (green) **(M)**. **F-G** Quantification of E. Estradiol increases proteins incorporated with puromycin (proteins^PURO^) along dendrites (**E**) and juxtaposing spine regions (**G**). n=13-15 cells from 3 independent replicates. **I-L** Quantification of H. RPS10 puncta density increased along dendrites (**I**) and juxtaposing spine regions (**J**). RPS10 puncta size is also increased along dendrites (**K**) and juxtaposing spine regions (**L**). n=12-15 cells from 3 independent replicates. **N-O** Quantification of M. Increased co-localisation between RPS10 and proteins^PURO^ RPS10 was observed along dendrites (**N**) and spine regions (**O**) in estradiol treated neurons. n=13- 15 cells from 3 independent replicates. Arrowheads denote localisation of puromycin/RPS10 at dendrites (yellow) and crude synaptic regions (white). Co-localisation between proteins is shown in white. Scale bar = 5µm **(E+H)**; 10 µm, 1µm (inset) **(M)**. Unpaired student’s t-tests, Welch’s correction. Error bars represent mean ± SEM; * p=<0.05, ** p = <0.01, *** p = <0.001.

To ascertain a more precise location of protein translation within neurons, instant Structured Illumination Microscopy (iSIM) super-resolution imaging was coupled with SUnSET-ICC. Elevated levels of proteins^PURO^ were detected in estradiol treated neurons along MAP2-positive dendrites (p=0.0017; **Figure 2E+F**) and areas juxtaposed to dendrites, which likely include synaptic regions (p=0.0002; **Figure 2E+G**). Examination of the distribution of the ribosomal protein S10 (RPS10), a component of the 40S ribosomal subunit, revealed that estradiol treatment increased RPS10 expression along and juxtaposed to dendrites (puncta per 10μm; dendrites: p=0.0159; crude synaptic regions: p=0.0238; **Figure 2H-J**). In addition, RPS10 puncta were larger in estradiol treated neurons compared to vehicle treated neurons (puncta area μm^2^; dendrites: p=0.0211; crude synaptic regions: p=0.0258; **Figure 2H-L**). Moreover, we observed an increase in the number of RPS10 puncta that co-localised with proteins^PURO^, indicating actively translating ribosomes, along and juxtaposed to dendrites in estradiol treated neurons (puncta per 10μm; dendrites: p=0.0205; crude synaptic regions: p=0.0393; **Figure 2M-O**). Taken together, these data suggest that estradiol increases the number of actively translating ribosomes along dendrites and near synapses, as indicated by an increase in the number of puromycylated polypeptides and increased ribosomal protein number and size.

### Estradiol increases dendritic spine density and nascent protein expression at synapses in a protein translation-dependent manner

Estradiol has been shown to induce rapid and long-lasting increases in dendritic spine density in male and female (OVX) hippocampal neurons (Mukai *et al*., 2007; Phan *et al*., 2015; Jacome *et al*., 2016; Tuscher *et al*., 2016; Luine *et al*., 2018). We thus, investigated whether this increase was protein synthesis dependent. Similar to what has been previously reported, estradiol treatment increased spine density after 2 hours (**Figure 3A+B**). Although estradiol did not have an effect on the average size of dendritic spines (**Figure 3C**), there was a shift in the number of spines that had a larger spine area in neurons treated with estradiol (**Figure 3D**). Interestingly, in neurons pre-treated with anisomycin, no difference in spine density was seen between vehicle and estradiol treated conditions (**Figure 3B+C**) suggesting that protein synthesis is required either for estadiol-induced spine formation or maintenance of nascently formed spines.

**Figure 3.**
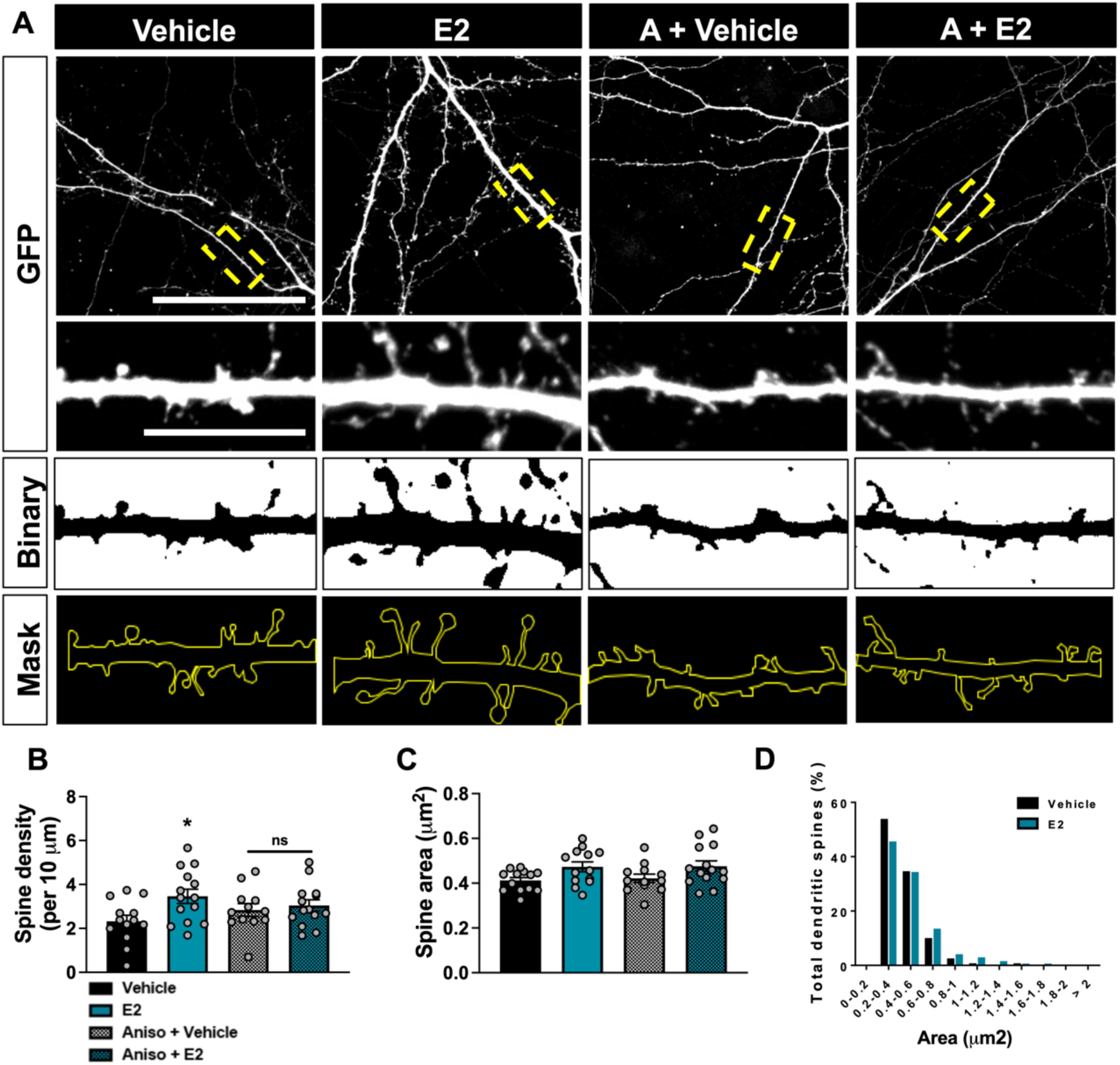
Estradiol-dependent protein synthesis is required for estradiol-mediated increases in dendritic spines. **A** Representative confocal images of DIV20 primary hippocampal neurons transfected with eGFP and treated with estradiol (E2; 2 hours) or vehicle (DMSO; 2 hours) with or without protein translation inhibitor, anisomycin (Aniso (A)). **B+C** Quantification of A. Estradiol increases dendritic spine number within 2 hours, which is blocked by anisomycin **(B**). Treatment had no effect on spine area **(C)**. **D** Histogram of spine area illustrated that estradiol increases dendritic spines with a larger area. n=12-14 cells from 4 independent replicates. Yellow dashed box denotes section of dendrite displayed in inset. Scale bar = 50µm, 10µm (inset). One-way ANOVA, Bonferroni corrected. Error bars represent mean ± SEM; * p=<0.05, ns = not significant.

As we had detected an increase in actively translating ribosomes along dendrites and near synapses, we reasoned that this would be also reflected by an increase in the presence of nascent protein in dendritic spines. To investigate this, fluorescent non-canonical amino acid tagging (FUNCAT) (Dieterich *et al*., 2010) was employed to metabolically tag and visualise newly synthesised proteins. Primary hippocampal neurons treated with estradiol for 2 hours exhibited an increase in the presence of azidohomoalanine (AHA)-tagged newly synthesised proteins (proteins^AHA^) along dendrites and within dendritic spines (dendrites: p=0.0002; spines: p=<0.0001; **Figure 4A-C**). The increase in proteins^AHA^ expression was attenuated by anisomycin in both dendrites (p=0.9999) and dendritic spines (p=0.9568; **Figure 4A- C**).

**Figure 4.**
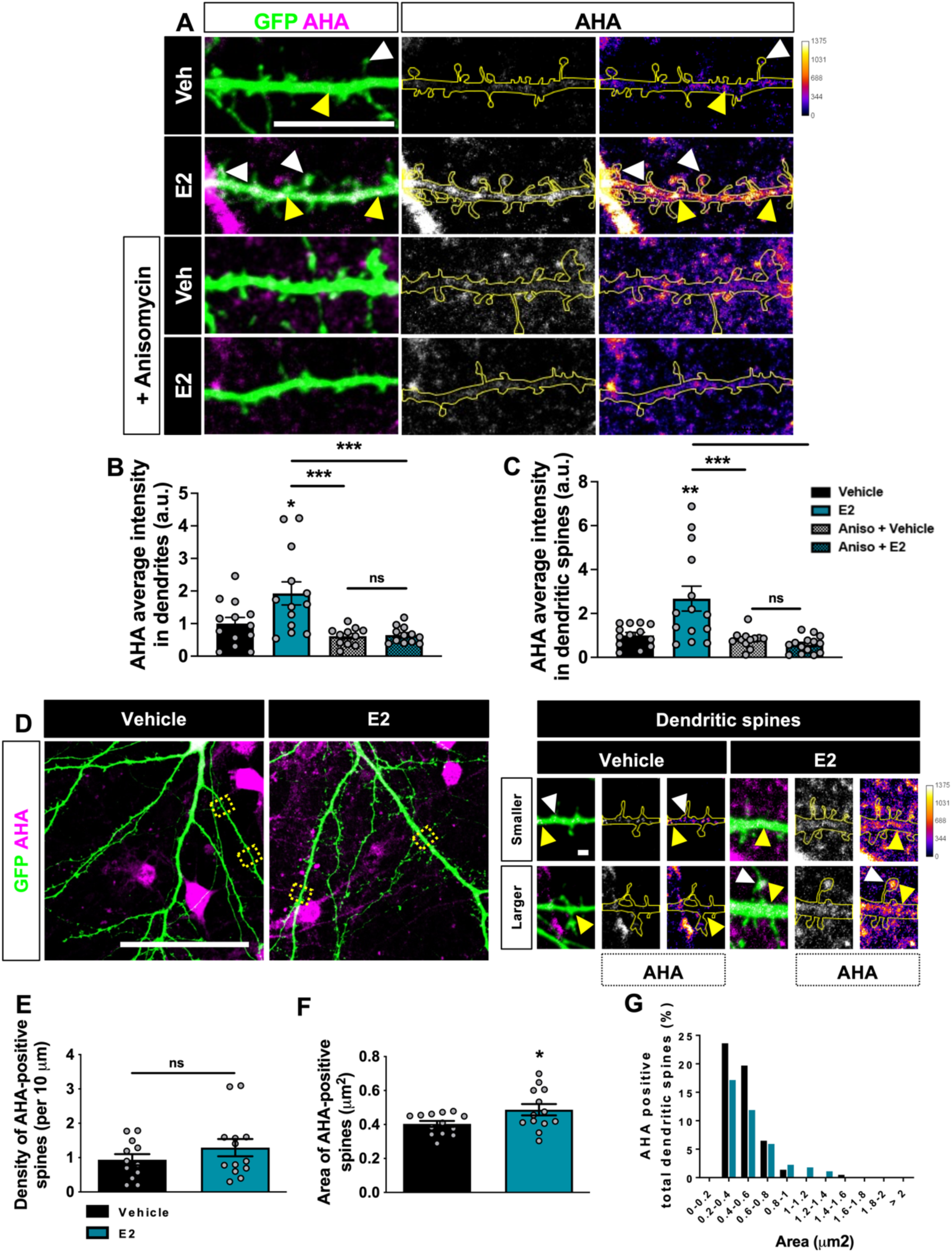
Newly synthesised proteins are increased within dendritic spines in estradiol treated neurons. **A+D** Representative confocal images of DIV20 primary hippocampal neurons transfected with eGFP. Neurons were then treated with vehicle control (DMSO) or estradiol (E2; 10nM) and azide-bearing azidohomoalanine (AHA; 4mM) for 2 hours. For some, neurons were pre-treated with protein translation inhibitor, anisomycin (Aniso, 40µM; 30 minutes). Following fixation, AHA was tagged with Alexa Fluor Alkyne-555 (2mM; overnight) and neurons were immunostained for GFP. **B+C** Quantification of A. Estradiol increases AHA-tagged newly synthesised proteins (proteins^AHA^), an effect prevented by pretreatment with anisomycin, within dendrites (**B**) and dendritic spines (**C**). n=11-13 cells from 4 independent replicates. **E+F** Quantification of D. The number of proteins^AHA^ -positive dendritic spines did not change with treatment (**E**). However, following treatment proteins^AHA^-positive dendritic spines had larger areas on average (**F**). **G** Histogram of spine area illustrating that following estradiol treatment a larger portion of proteins^AHA^-positive dendritic spines have increased area. n=12-13 cells from 4 independent replicates. Arrowheads denote localisation of proteins^AHA^ at dendrites (yellow) and within dendritic spines (white). Co-localisation between proteins is shown in white. Yellow dashed box denotes section of dendrite displayed in inset. Scale bar 10µm **(A)**; 50µm, 10µm (inset) **(D)**. One-way ANOVA, Bonferroni corrected. Error bars represent mean ± SEM; * p=<0.05, *** p=<0.001, ns = not significant.

We further examined if a sub-set of spines were enriched for nascent proteins following treatment. Estradiol treated neurons showed no significant difference in the number of AHA-positive spines compared to vehicle treated neurons (p=0.2572; **Figure 4D+E**). However, analysis of spine morphology revealed that in estradiol treated neurons, spines containing proteins^AHA^ were on average larger compared to those in vehicle treated cells (p=0.0400; **Figure 4D+F**). Consistent with this observation, categorising AHA-positive spines by spine area revealed that spines containing proteins^AHA^ following estradiol treatment were shifted towards those with larger spine areas. Estradiol increased AHA intensity in larger spines, with an area larger than 0.8µm², where more AHA could be observed in smaller spines, areas smaller than 0.8µm², in vehicle treated cells (**Figure 4.D+G**). This suggests an accumulation of new proteins in larger spines following estradiol treatment. Taken together, these data provide supporting evidence that estradiol-mediated increases in local protein synthesis promotes an elevation in newly formed dendritic spines as well as newly synthesised proteins along dendrites and within dendritic spines.

### Estradiol regulates the expression of the synaptic proteome in male and female hippocampus

Given that estradiol rapidly increases protein synthesis in both male and OVX female hippocampus, we were interested in establishing how estradiol was affecting the synaptic proteome. Male and OVX female hippocampal slices were treated for 2 hours with estradiol or vehicle and the expression of candidate pre- and post-synaptic proteins (**Figure 5A+B**). In male hippocampal slices, estradiol treatment significantly increased expression of both scaffolding protein post synaptic density 95 (PSD-95) (p<0.0001) and the AMPAR subunit GluA1 (p=0.0008) (**Figure 5B+C**). No significant change in the NMDAR subunit GluN1 expression levels was observed following treatment (p=0.6470); but estradiol induced an increase expression levels of both subunits GluN2A (p=0.0018) and GluN2B (p=0.0079) within the same timeframe (**Figure 5B+C**). Treatment with estradiol in OVX female hippocampal slices resulted in an increase in PSD-95 (p=0.0003) and GluA1 (p<0.0001) expression, as seen in male hippocampal slices. Intriguingly, GluN1 expression was also increased following treatment (p=0.0202; **Figure 5B+D**). Both GluN2A (p=0.0007) and GluN2B (p=0.0204) expression were both increased in response to estradiol treatment (**Figure 5B+D**).

**Figure 5.**
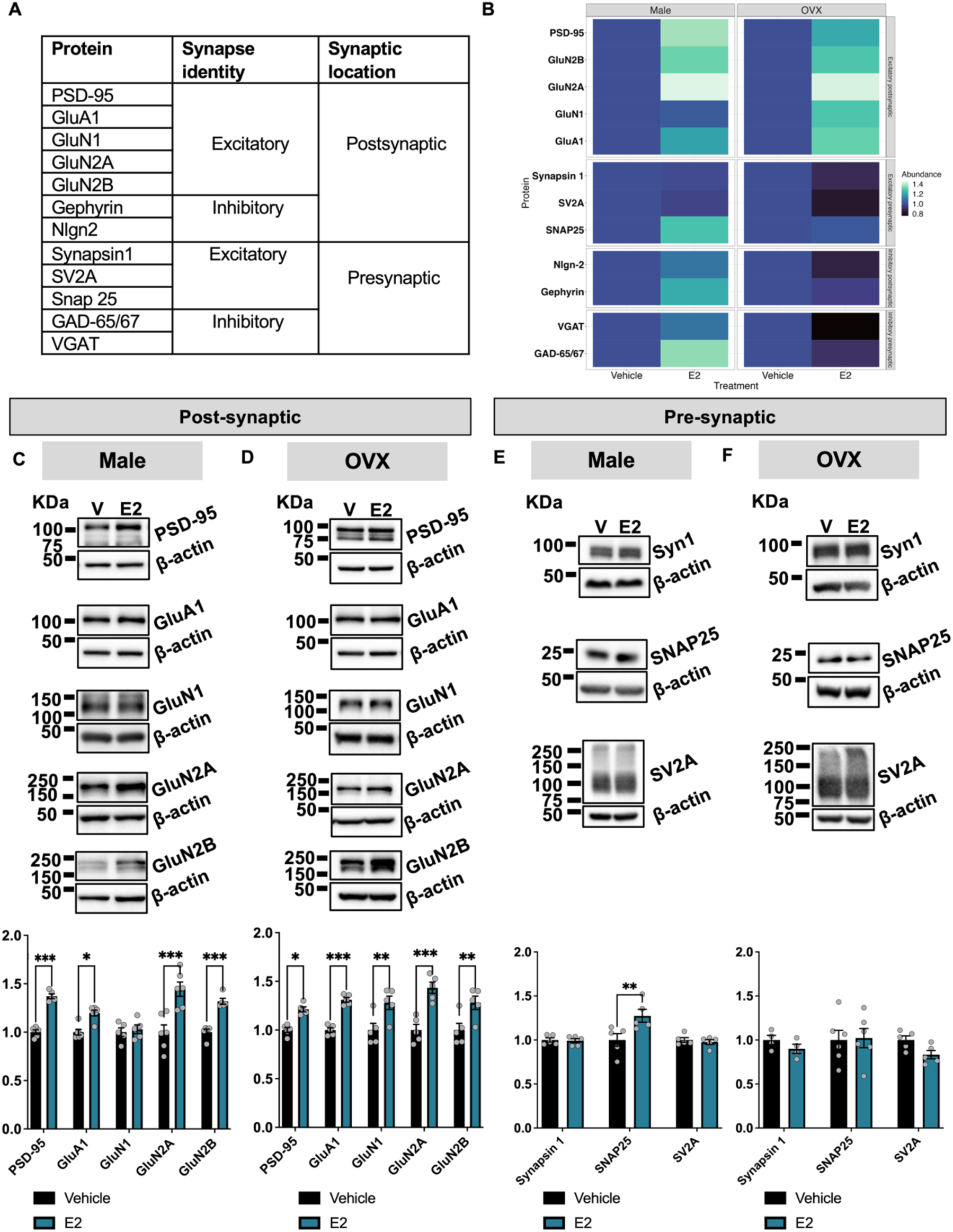
The excitatory synaptic proteome is regulated differentially in males and OVX females following estradiol exposure. **A** List of synaptic proteins assessed by western blotting following estradiol treatment of acute hippocampal slices from 10-12 week old male or and ovariectomised (OVX) mice. **B** Heat map demonstrating the impact of estradiol treatment on synaptic protein expression. **C-F** Representative western blots and quantification of male or OXV acute hippocampal slices treated with vehicle (DMSO; V) or estradiol (E2; 10nM) for 2 hours within the same animal. Samples were immunoblotted for scaffolding protein post synaptic density 95 (PSD-95), AMPAR subunit GluA1, the NMDA receptor subunits GluN1, GluN2A and GluN2B, the neurotransmitter release regulator synapsin 1 (Syn1), synaptosomal associated protein 25 (SNAP25) and synaptic vesicle protein 2A (SV2A) and normalised to housekeeper, β-actin. **C** In the male hippocampus, estradiol increased the expression of PSD-95, GluA1, GluN2A and GluN2B but not GluN1. **D** In OVX female hippocampus, estradiol increased the expression of PSD-95, GluA1, GluN1, GluN2A and GluN2B. **E** In the male hippocampus, estradiol had no effect on Syn1 or SV2A, but increases SNAP-25 expression. **F** In the OVX female hippocampus, estradiol had no effect on Syn1 and SNAP-25, but decreased SV2A expression. Male: n=5 animals for PSD-95, GluN1, GluN2B, synapsin1, SNAP25 and SV2A/condition; n=6 animals for GluA1 and GluN2A/condition; OVX female: n=4 animals for synapsin1/condition; n=5 for PSD-95, GluA1, GluN2A and GluN2B; n=6 animals for GluN1 and SNAP25/condition. Unpaired student’s t-tests, Welch’s correction. Mann-Whitney test was used with non-parametric data sets. Error bars represent mean ± SEM; * p=<0.05, ** p = <0.01, *** p = <0.001, *** p = <0.001, ns = not significant.

As estradiol has also been shown to regulate inhibitory synapses (Mukherjee *et al*., 2017), we next assessed the expression of synaptic proteins restricted to inhibitory post-synaptic sites. Estradiol significantly increased expression of the post- synaptic scaffolding protein gephyrin in the male (p<0.0001; **Figure 5B + S2A**) but not in OVX female hippocampal slices (p=0.7134; **Figure 5B + S2B**). Conversely, no significant difference in adhesion protein neuroligin-2 (Nlgn2) expression was found in either male (p=0.1096; **Figure 5B + S2C**) or OVX female (p=0.2104; **Figure 5B + S2D**) hippocampal slices. Taken together, these data suggest that estradiol rapidly increases the expression of synaptic proteins that are highly abundant at the PSD of excitatory synapses in both the male and OVX female mice hippocampus but has sex-specific effects on the expression of inhibitory post-synaptic proteins.

### Estradiol differentially regulates the excitatory pre-synaptic proteome in a sex dependent manner

To gain a more complete understanding of how estradiol may be impacting synapses, we next assessed the expression of pre-synaptic proteins abundant at either excitatory or inhibitory synapses. Estradiol treatment did not alter expression levels of the neurotransmitter release regulator synapsin 1 (Syn1) in either males (p=0.8082; **Figure 5E**) or OVX female (p=0.2000; **Figure 5F**) hippocampal slices. However, sex-specific differences in the expression of both synaptosomal associated protein 25 (SNAP25) and synaptic vesicle protein 2A (SV2A) were observed. Estradiol treatment increased SNAP25 expression in males (p=0.0280; **Figure 5E**), but not OVX female hippocampal slices (p=0.8910; **Figure 5F**). Conversely, estradiol had no effect on SV2A expression in the male hippocampal slices (p=0.5485; **Figure 5E**) but decreased expression of this pre-synaptic protein in OVX female hippocampal slices (p=0.0357; **Figure 5F**).

When we examined proteins predominately restricted to inhibitory pre-synaptic sites, we found that estradiol treatment significantly increased glutamic acid decarboxylase 65 and 67 (GAD-65/67) expression in the male (p=0.0286; **Figure S2E**) but not in OVX female (p=0.3429; **Figure S2F**) hippocampal slices. Interestingly, estradiol did not alter the expression of vesicular GABA transporter (VGAT) in male (p=0.3351; **Figure S2G**), but significantly reduced its expression in OVX female (p=0.0036; **Figure S2H**) slices. Taken together, these data demonstrate that estradiol has sex-specific effects on the expression of protein localised to either excitatory or inhibitory pre-synaptic terminals. Moreover, these proteins displayed both increased and decreased expression in response to treatment, indicating that estradiol may be regulating proteostasis at synapses in general.

### Estradiol increases PSD-95 and GluN2B in a local protein synthesis manner

We next wanted to investigate whether estradiol was increasing the expression of select synaptic proteins via a local protein synthesis mechanism. To this end, we focused on PSD-95 and GluN2B as both proteins increased in expression following estradiol treatment in male and female hippocampal slices and have been implicated in estradiol’s effects on synaptic function (Akama & McEwen, 2003; Smith & McMahon, 2006; Liu *et al*., 2008). As expected, pre-treatment with anisomycin attenuated estradiol-mediated increases in PSD-95 (p=0.0022) and GluN2B (p=0.0003) in male hippocampal slices (**Figure 6A+B**). Comparably, a similar consequence was observed in OVX female hippocampal slices. Estradiol-dependent increases in PSD-95 (p=0.0007) and GluN2B (p=0.0016) expression were blocked by pre-treatment with anisomycin (**Figure 6C+D**). Together, these data are consistent with estradiol driving an increase in expression of these proteins via a mCap- translation mechanism.

**Figure 6.**
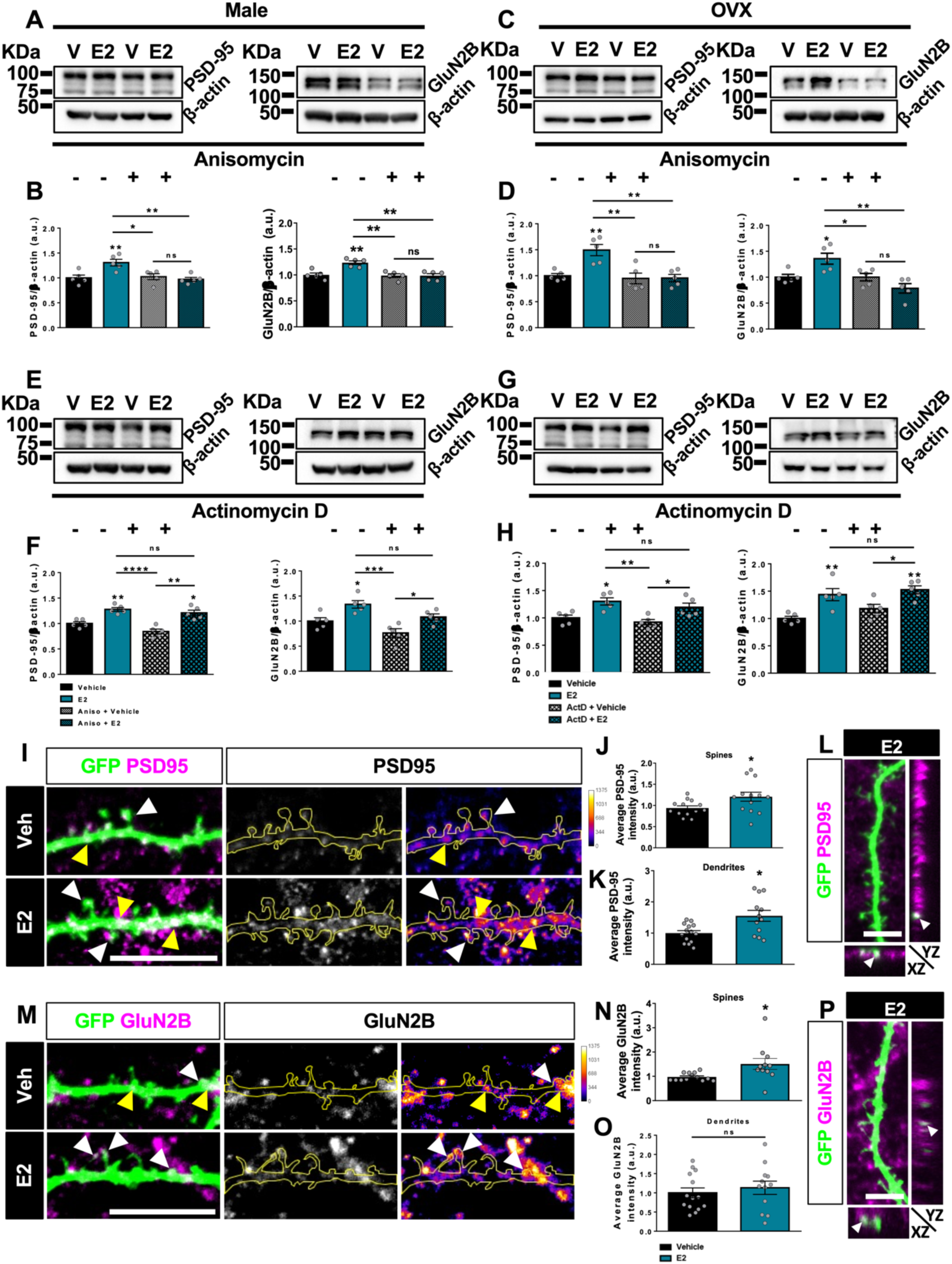
Estradiol increases PSD-95 and GluN2B in a local protein synthesis dependent manner. **A-H** Representative western blots and quantification of acute hippocampal slices from 10-12 week old male **(A+E)** and ovariectomised (OVX) mice **(C+G)** pre-treated for 30 minutes with no inhibitor, protein translation inhibitor, anisomycin (Aniso, 40µM) or RNA polymerase II inhibitor, actinomycin D (ActD; 25µM) within the same animal. Slices were processed for western blotting, immunoblotted for PSD-95 and GluN2B and normalised to housekeeper, β-actin. Anisomycin inhibited estradiol-mediated increases of **(B)** PSD-95 and GluN2B in the male hippocampus. Anisomycin also inhibited estradiol-mediated increases of **(D)** PSD-95 and GluN2B in the OVX females. Whereas, estradiol continued to mediate an increase **(F)** PSD-95 and GluN2B expression in males. This was also recapitulated in both **(H)** PSD-95 and GluN2B in the OVX female hippocampus. n=5 per animal for both male and OVX female/condition). **I, L, M, P** Representative confocal images (60x) and quantification of DIV20 primary hippocampal neurons transfected with eGFP and treated with vehicle control (DMSO; V; 2 hours) or estradiol (E2; 10nM; 2 hours). Neurons were fixed and immunostained for GFP (green) and PSD-95 (magenta) **(I+L)** or GluN2B (magenta) **(M+P)**. **J+K** Quantification of I. PSD-95 is increased within dendritic spines (**J**) and along dendrites (**K**) in estradiol neurons. n=14 for vehicle, 13 **(J)** and 12 **(K)** for estradiol cells over 4 different biological replicates. **L** Orthogonal projections confirmed localisation of PSD-95 within spines in estradiol treated neurons. **N+O** Quantification of M. GluN2B is increased within dendritic spines (**N**) but not extrasynaptically along dendrites (**O**) in estradiol treated neurons. **P** Orthogonal projections confirmed localisation of GluN2B within dendritic spines in estradiol treated neurons. n=13 **(N)**, 14 **(O)** for vehicle, 12 for estradiol cells over 4 different biological replicates. Arrowheads denote localisation of PSD-95/GluN2B at dendrites (yellow) and within crude spine regions (white). Co-localisation between proteins is shown in white. Scale bar = 10µm **(I+M)**; 5µm **(L+P)**. Unpaired student’s t-tests, Welch’s correction; One-way ANOVA, Bonferroni corrected. Error bars represent mean ± SEM; * p=<0.05, ** p = <0.01, *** p = <0.001, *** p = <0.001, ns = not significant.

We next investigated whether estradiol-mediated increases were maintained or blocked in the presence of the gene transcription inhibitor actinomycin D (ActD). In male hippocampal slices, estradiol was still able to increase the expression of PSD- 95 and GluN2B in the presence of ActD (PSD-95, p<0.0001; GluN2B, p=0.0005; **Figure 6E+F**). Similarly, in OVX female hippocampal slices - estradiol ActD did not attenuate estradiol’s ability to increase PSD-95 or GluN2B expression levels (PSD95, p=0.0019; GluN2B, p=0.0010; **Figure 6G+H**). These data suggest that the estradiol-mediated increases in PSD-95 and GluN2B protein expression levels is independent of gene transcription. Consistent with this, assessment of both synaptic proteins at the mRNA level revealed that estradiol induced no significant change in either *dlg4* (*dlg4*, p=0.7584; **Figure S4A**) or *grin2b* (*grin2b*, p=0.9496; **Figure S4B**) expression in primary hippocampal neurons. Taken together, these data provide evidence that estradiol increases the expression of synaptic proteins PSD-95 and GluN2B through a gene transcription-independent manner.

We next sought to investigate the sub-cellular location of PSD-95 and GluN2B expression following estradiol treatment. Estradiol increased PSD-95 (**Figure S4C**) and GluN2B (**Figure S4D**) expression in crude synaptosomal fractions generated from primary hippocampal cultures. Using immunocytochemistry, increased PSD-95 expression was observed within spines (p=0.0367) and dendrites (p=0.0108) (**Figure 6I-K**) in primary hippocampal neurons. Specifically, PSD-95 could be visualised within spine heads of estradiol treated neurons (**Figure 6L**). Estradiol also increased GluN2B expression at spines (p=0.0403); no change in GluN2B expression was detected along dendrites (p=0.5438; **Figure 6M-P**). Collectively, these data suggest that after 2 hour treatment with estradiol, PSD-95 and GluN2B expression was increased within dendritic spines. As estradiol has no significant effect on their respective mRNA levels, these results suggest that estradiol may be engaging dendritic mRNA to locally translate these proteins.

### mTOR mediates estradiol-induced protein synthesis in a sex-specific manner

mTOR kinase signalling has been implicated in mediating local protein synthesis (Costa-Mattioli *et al*., 2009; Hoeffer & Klann, 2010) and multiple studies have demonstrated estradiol’s ability to rapidly phosphorylate mTOR in the hippocampus and cortex (Fortress *et al*., 2013; Briz & Baudry, 2014; Sellers *et al*., 2015b). As mTOR is upstream of several protein translation initiation elements, we reasoned that estradiol-induced protein synthesis may be occurring through a mTOR-dependent mechanism. In agreement with this idea, we found that estradiol increased mTOR activation in a time-dependent manner in mixed sex primary neuronal cultures (p=0.0018; **Figure S5A-B**), similar to previous reports (Briz & Baudry, 2014; Sellers *et al*., 2015b). mTOR activation was significantly increased at 30 minutes (p=0.0012) and remained increased at 1 hour (p=0.0128). However, 2 hours after treatment, mTOR phosphorylation decreased but remained above baseline. These data indicated that estradiol activates mTOR in a time-dependent manner, with maximal activation observed prior to estradiol-induced protein synthesis.

We next sought to determine whether mTOR was required for estradiol-induced increased global protein synthesis. Male and OVX female hippocampal slices were pre-treated with the mTOR inhibitor, rapamycin (1 µM), 30 minutes prior to estradiol or vehicle treatment; puromycin was subsequently added to the slices and proteins^PURO^ quantified. As seen before, estradiol alone increased protein synthesis in male and OXV female hippocampal slices. Interestingly, pre-treatment with rapamycin inhibited estradiol-mediated increase in protein synthesis in male hippocampal slices (p=0.0004; **Figure 7A+C**). However, in OVX female hippocampal slices, pre-treatment with rapamycin had no effect on estradiol-induced protein synthesis (p=0.0043; **Figure 7B+D**). Together, these data indicate that mTOR was required for estradiol-induced global protein synthesis in male hippocampal slices only, and thus was part of a sex-specific mechanism.

**Figure 7.**
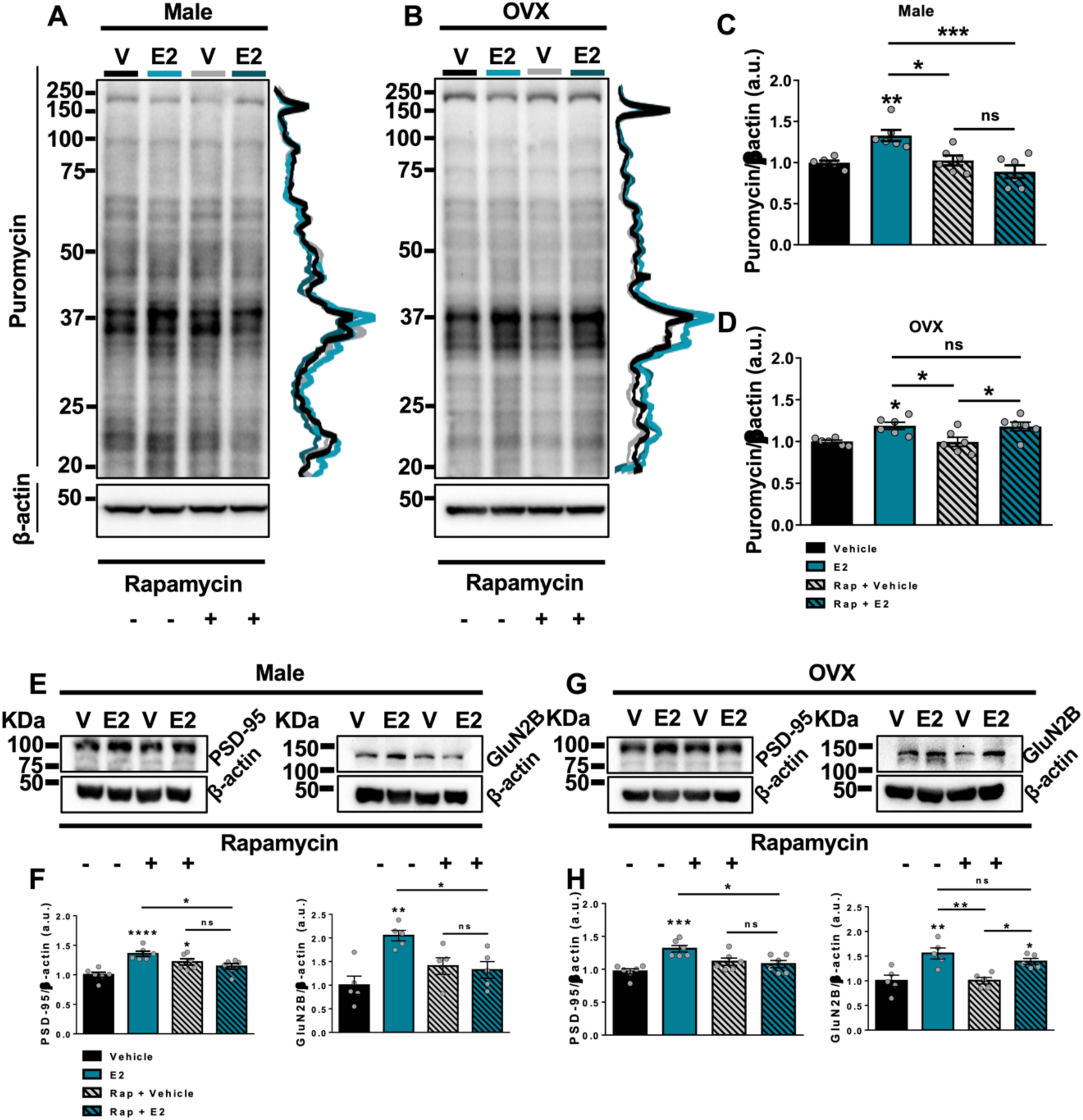
mTOR is required for estradiol to increase global protein synthesis in the male, but not the OVX female, hippocampus. **A, B, E, G** Representative western blots and quantification of acute hippocampal slices from 10-12 week old male **(A+E)** and ovariectomised (OVX) mice **(B+G)** pre-treated with mTOR signalling pathway inhibitor, rapamycin (Rap; 1µM; 30 minutes) or no inhibitor followed by estradiol (E2; 10nM; 2 hours) or vehicle control (DMSO; V; 2 hours) **(E+G)** and puromycin (SUnSET assay, 5µg/mL, last 30 minutes) **(A+B)** treatment within the same animal. Slices were processed for western blotting, immunoblotted for puromycin and normalised to housekeeper, β-actin. **C+D** Quantification of A+B. Estradiol-mediated increases in the rate of protein synthesis were inhibited in the male (**C**) but not the OVX female (**D**) hippocampus. **F+H** Quantification of E and G, respectively. Rapamycin inhibited estradiol-mediated increases of **(F)** PSD-95 and GluN2B. In OVX females, rapamycin inhibited estradiol-mediated increase of **(H)** PSD-95 but not GluN2B. Line-scan next to each western blot is a visual representation of intensity changes within puromycin bands. Male: n=6 animals for both male and OVX female/condition across all experiments. One-way ANOVA, Bonferroni corrected. Error bars represent mean±SEM; * p=<0.05, ** p = <0.01, *** p = <0.001, *** p = <0.001, **** p = <0.0001, ns = not significant.

### mTOR is required for estradiol-dependent increase of specific synaptic proteins in the male and OVX female hippocampal slices

Although mTOR may be important for global protein synthesis in males, it is unclear if this signalling pathway is needed for regulating synaptic protein expression. Hence, we tested PSD-95 and GluN2B expression levels in acute hippocampal slices upon estradiol in combination with rapamycin treatment. Consistent with what we observed when examining global protein synthesis, rapamycin blocked estradiol-mediated increases in PSD-95 (p=0.0003) and GluN2B (p=0.0033) expression in male hippocampal slices (**Figure 7E+F**). A post hoc analysis confirmed an attenuation of PSD-95 (p>0.9999) and GluN2B expression (p>0.9999) in males.

Interestingly, when assessed in the OVX female hippocampal slices, rapamycin inhibition of mTOR signalling blocked the increase of PSD-95 (p=0.0009) but not GluN2B (p=0.0009) (**Figure 7G+H**). Post-hoc analysis confirmed that rapamycin only attenuated estradiol-mediated increases in PSD-95 (p>0.9999) but not GluN2B (p=0.0390) expression in OVX females. This suggested that mTOR was not required for estradiol’s effect on global protein synthesis and increased GluN2B expression in OVX female hippocampus, it is required, however, for estradiol-mediated increases in PSD-95 expression.

## Discussion

Estradiol has been consistently reported to increase the expression of a number of synaptic proteins which have been linked with increased synaptic function and effects on cognition (Smith & McMahon, 2006; Jelks *et al*., 2007; Liu *et al*., 2008; Smith *et al*., 2009; Vedder *et al*., 2013; Phan *et al*., 2015; Avila *et al*., 2017). However, the mechanism by which estradiol increases the expression of synaptic proteins remains unclear. In this study, we have directly tested whether estradiol regulates the rate of protein synthesis in hippocampal neurons, and if estradiol-mediated increases in synaptic proteins occurs via a local protein synthesis manner. We further have tested the requirement of the mTOR pathway in mediating this mechanism, and overall, have assessed whether any of estradiol’s effects on protein translation occur in a sex specific manner. We find that: **(i)** estradiol increases global protein synthesis in a gene-transcription independent manner in both male and female hippocampus within 2 hours; **(ii)** estradiol-modulation of protein synthesis is accompanied by an increase in the number of actively translating ribosomes and newly synthesised proteins along dendrites and at synapses; **(iii)** estradiol increases the expression of specific excitatory and inhibitory post-synaptic proteins in a sex-specific and local protein synthesis manner, and decreases the expression of specific pre-synaptic proteins in a sex-specific manner implicating a more general role for estradiol in regulating proteostasis; **(iv)** and that the requirement of the mTOR pathway for estradiol- modulation of local protein synthesis and regulation of synaptic protein expression also occurs in a sex-specific manner, with a more pronounced effect in the male hippocampus compared to the female.

The finding that estradiol can modulate protein synthesis in male and female hippocampal slices within a rapid time frame (2 hours) is consistent with observation from previous studies. Estradiol has previously been shown to regulate the activity of kinases involved in regulating protein translation, such as 4EBP1, p70S6K and RPS6 within a similar time frame (Akama & McEwen, 2003; Sarkar *et al*., 2010; Fortress *et al*., 2013; Briz & Baudry, 2014). While these studies have suggested that estradiol can regulate protein synthesis in neurons, they have not directly assessed protein synthesis or investigated whether the effects occur independently of gene transcription. Using the SUnSET approach to tag elongating polypeptide chains, we have been able to demonstrate that estradiol causes a time-dependent increase in the rate of protein synthesis in both male and female hippocampal slices. Moreover, we find that estradiol-induced increase in protein synthesis occurs even in the presence of transcription inhibition, and can be observed in crude synaptic preparations, indicating that estradiol is regulating a local protein synthesis mechanism in both male and female hippocampal slices.

Owing to the nature of hippocampal slice preparations, it is not possible to determine whether estradiol-mediated increases in protein synthesis are occurring in neurons, glia cells or both. Indeed, whilst SUnSET provides information about the rate (SUnSET-WB) of translation, SUnSET-ICC gives information of where actively translating ribosomes are located. In addition, FUNCAT shows newly synthesised proteins. Thus, combining both techniques provides information about the amount of translation and where nascent proteins are targeted following a pharmacological treatment. Using this combination of approaches, we observed an increase in proteins^PURO^ along and juxtaposed to MAP2-positive dendrites of primary hippocampal neurons. As we designed SUnSET-ICC with short puromycin incubation times, this suggests that estradiol regulates local protein synthesis in neurons, and moreover near synaptic sites. This is consistent with the findings by Sarkar et al., who reported that estradiol rapidly induced dendritic translation, assessed by examining CAMKIIα 3’-UTR driven GFP expression in primary neurons (Sarkar *et al*., 2010). Interestingly, fluctuations of estradiol levels during the rat estrous cycle has also been shown to trigger polyribosome accumulation at dendrites (McCarthy & Milner, 2003). In line with this observation, we found an increase in the number and size of RPS10 along dendrites in response to estradiol treatment. This was accompanied by an increased number of colocalised RPS10 and proteins^PURO^ puncta along dendrites. As increased RPS10 puncta was also observed near putative synaptic regions, it raised the possibility that estradiol was engaging translational machinery along dendrites to promote the synthesis of new proteins within the vicinity of synapses. Consistent with this idea, we observed an increase in nascently synthesised proteins (proteins^AHA^) along dendrites and specifically in dendritic spines. Interestingly, we also found that estradiol increased dendritic spine density and a shift towards larger spines after 2 hours, which is in line with previous findings (Phan *et al*., 2015; Jacome *et al*., 2016; Tuscher *et al*., 2016; Avila *et al*., 2017). Concurrent with this, an increase in the amount of proteins^AHA^ within dendritic spines was observed and blocking protein synthesis attenuated estradiol’s effects on dendritic spine density. However, it is not possible to determine whether the synthesis of nascent proteins is required for estradiol-induced spine formation or the maintenance of remodelled spines. Of note, BDNF-mediated spine enlargement has been shown to be dependent on protein synthesis (Tanaka *et al*., 2008). Conversely, estradiol-induced spine formation in hippocampal neurons is not dependent on protein synthesis (Sheppard *et al*., 2021). Thus, it is possible that in our experiments, synthesis of nascent protein is required for the maintenance, but not the formation, of estradiol-induced nascent spines. Taken together, these data illustrate that by regulating local protein synthesis along dendrites, estradiol can rapidly and locally synthesis and target new proteins to synapses.

Multiple studies have shown that estradiol can modulate the expression of synaptic proteins within a rapid time frame in both male and female hippocampus (Akama & McEwen, 2003; Smith & McMahon, 2006; Jelks *et al*., 2007; Liu *et al*., 2008; Avila *et al*., 2017). We assessed the impact of estradiol on the expression of a range of pre- and post-synaptic proteins, predominately found at either excitatory or inhibitory synapses, in male and female hippocampi. This has revealed a sex-specific regulation of both excitatory and inhibitory post-synaptic protein expression by estradiol. Further, it illustrated that estradiol was able to regulate pre-synaptic protein expression, also in a sex-specific manner. Overall, estradiol was seen to increase the expression of all excitatory post-synaptic proteins examined in both sexes, aside from GluN1, which was only seen to show increased expression in female hippocampal slices following treatment. Of the inhibitory post-synaptic proteins examined, only Gephyrin showed an increase in synaptic protein expression, and only in male hippocampal slices. Interestingly, when pre-synaptic proteins were examined, a sex-specific decrease in protein expression was observed. In male hippocampal slices Snap25 and GAD65/67 showed decreased expression, whereas Synapsin1 and VGAT1 were decreased in female hippocampal slices following treatment with estradiol. Not only do these results show that estradiol has a sex-specific effect on the examined pre-synaptic proteins, but further indicates that estradiol can also induce protein degradation. Interestingly, behavioural training has recently been shown to induce sex-specific changes in the ubiquitin proteasome system (Beamish *et al*., 2022). Moreover, estradiol has also been posited to also engage with the ubiquitin proteasome system to regulate protein degradation (Erli *et al*., 2020; Beamish & Frick, 2021). Overall, our observations that estradiol can regulate the expression of specific pre- and post-synaptic proteins within a rapid time frame are consistent with the effects of this steroid on synaptic plasticity (Liu *et al*., 2008; Kramar *et al*., 2013; Vedder *et al*., 2013; Jain & Woolley, 2023). Critically, these data highlight the importance of considering sex when assessing synaptic phenotypes. Indeed, a recent study examining the induction of LTP pre or post adolescent male or female rats revealed a sex-specific mechanism that was driven in part by the estrogen alpha receptor (Le *et al*., 2022). Whether the observed effects shown in this study contribute to these differences could be investigated in the future.

We focused on the impact of estradiol-induced increased expression of GluN2B and PSD-95, as both proteins have been implicated in mediating the effects of this steroid on synaptic plasticity and behaviour (Akama & McEwen, 2003; Smith & McMahon, 2006; Liu *et al*., 2008; Avila *et al*., 2017). Estradiol has been reported to increase the expression of PSD-95 located at synapses (Akama & McEwen, 2003; Liu *et al*., 2008; Srivastava *et al*., 2010; Sellers *et al*., 2015b), and are thought to contribute to the increased targeting of AMPA receptors to synapses, and thus drive estradiol- facilitation of synaptic plasticity (Srivastava *et al*., 2008; Phan *et al*., 2015; Sheppard *et al*., 2023). Multiple studies have also reported that GluN2B-containing NMDARs are critical for estradiol-induced enhancement in LTP (Smith & McMahon, 2006; Snyder *et al*., 2011; Vedder *et al*., 2013; Smith *et al*., 2016). Potier et al. recently reported that estradiol transiently decreased GluN2B surface diffusion and contribute to estradiol-mediated increase in dendritic spines (Potier *et al*., 2016). Consistent with such studies, and the observed increase in proteins^AHA^ at dendritic spines in this study, we observed an increase of both GluN2B and PSD-95 at excitatory synapses. Importantly, estradiol-driven increased expression of both proteins was not dependent on gene transcription, and thus consistent with a local protein synthesis mechanism. Perhaps one of the most striking findings we have observed in this study is the sex-specific contribution of mTOR to the observed phenotypes. mTOR was shown to be required to induce estradiol-dependent global protein synthesis in the male but not female hippocampus. Furthermore, mTOR was also required for estradiol-induced increase in PSD-95 and GluN2B expression levels in the male hippocampus and interestingly, only PSD-95 in OVX female hippocampus. As SUnSET is a measure of global protein synthesis, it is not possible to differentiate between potential contribution of multiple signalling pathways involved in mediating estradiol-mediated protein synthesis. Therefore, mTOR may be required for mediating a proportion of estradiol-mediated protein synthesis, which cannot be detected using the SUnSET assay. Consistent with this idea, we found that while inhibition of mTOR signalling was able to block estradiol-induced increased expression of GluN2B in male but not female hippocampus, it was sufficient to block the effect of estradiol on the expression of PSD-95 in both male and female hippocampal slices. This suggests that mTOR has a greater role in mediating the effects of estradiol on mediating local protein synthesis of synaptic proteins in male hippocampal slices, this pathway play a more subtle contribution to these effects in female hippocampal neurons.

In summary, this study supports the idea that estradiol rapidly regulates the expression of synaptic proteins through local protein synthesis. The rapid increase in expression of synaptic proteins could lead to lasting alterations in synaptic function and contribute to estradiol-induced changes in cognitive function. At synapses, proteins are continually synthesized and degraded to modulate cellular function in response to various stimuli, such as action potentials, hormones, or external factors (Alvarez-Castelao & Schuman, 2015). Interestingly, we find that estradiol has a bidirectional influence on the expression of specific synaptic proteins, suggesting a broader role in regulating proteostasis. Notably, estradiol utilizes the mTOR signaling pathway in a sex-specific manner to control local protein synthesis of synaptic proteins. This finding contributes to our understanding of how estradiol impacts synaptic plasticity, and is consistent with the concept of latent (or hidden) effects, as discussed by Oberlander and Woolley (2016), where different molecular mechanisms may lead to similar outcomes in both sexes. It also emphasizes the importance of accounting for sex when investigating the molecular mechanisms underlying cellular processes. Moreover, given the presence of sex differences in various psychiatric disorders and neurodegenerative diseases, incorporating sex as a variable is crucial for tailoring treatments to each gender. Indeed, as there is mounting evidence of the therapeutic potential of estrogens or estrogen-based compounds, future studies exploring their therapeutic applications should carefully consider sex as a variable.

## Methods

### Animals

Mixed sex hippocampal cultures were prepared from Sprague-Dawley rat E18 embryos (Charles River Laboratories, U.K.) as previously described (Srivastava *et al*., 2008). The experimental procedures were carried out in accordance with the Home Office Animals (Scientific procedures) Act, United Kingdom, 1986. All acute cortical and hippocampal slices were prepared from intact male and ovariectomised (OVX) female C57BL/6J mice (The Jackson Laboratory, Maine, U.S.A) between the ages of 10-12 weeks. The female mice were OVX-ed by the vendor at 10 weeks and transported after 1 week of recovery; intact male mice were transported at 10 weeks. All mice were habituated for 3 days before experimental procedures were carried out. Age of ovariectomy and time of experiment post ovariectomy and transportation of the female mice was kept constant to minimise any confounding effects.

### Acute slice preparation

Mice were anaesthetised with isoflurane, followed by decapitation. Brain tissue was then rapidly removed and sliced 350 µm thick coronally in carbo-oxygenated (95% oxygen, 5% carbon dioxide), ice cold cutting solution using a Leica VT1000 vibratome. Hippocampi were rapidly dissected from the slices and maintained in a recovery chamber with carbo-oxygenated Ringer’s solution for 1 hour at 32°C. Slices were then transferred into 6-well plates containing 5-10 mL of freshly carbo-oxygenated Ringer’s solution in the presence of various drugs. Slices were collected and frozen at -80°C until lysing.

### Pharmacological treatments

Primary neurons and acute slices were treated with the following drugs: 17β-estradiol (estradiol; E2; 10nM; Sigma: E8875), anisomycin (Aniso; 40µM; Sigma: A9789), actinomycin D (ActD; 25µM; Sigma: A4262) and rapamycin (Rap; 1µM; Cell Signalling: 9904). All pharmacological treatments were carried out in artificial cerebral spinal fluid (aCSF; in mM: 125 NaCl; 2.5 KCl; 26.2 NaHCO_3_; 1 NaH_2_PO_4_; 11 glucose; 5 HEPES pH 7.4; 2.5 CaCl_2_; 1.25 MgCl_2_) in primary neurons. and Ringer’s solution (in mM: 126 NaCl; 10 glucose; 2 MgCl_2_; 2 CaCl_2_; 2.5 KCl; 1.25 NaH_2_PO_4_; 1.5 mM C_3_H_3_NaO_3_; 1 L-glutamine; 2.6 NaHCO_3_ for acute slices. Neurons were pre-treated in aCSF and slices were recovered in for 1 hour prior to application of the drug compounds. If inhibitors were relevant to the experiment, neurons were in aCSF for 30 minutes after which the inhibitors would be added for 30 minutes followed by pharmacological treatment; the inhibitors were added after 1 hour recovery for 30 minutes for acute slices. All compounds were diluted in dimethyl sulfoxide (DMSO). Vehicle controls were made up of solvent (DMSO) lacking the compounds and diluted as test compounds; the solvent was diluted to at least 0.1%.

### Surface sensing of Translation (SUnSET)

Puromycin treatments were added during the last 10 (neurons) or 30 minutes (slices) of pharmacological treatments at the final concentrations of 10 and 5 µg/ml respectively. The neurons or slices were then lysed for biochemistry or PFA-sucrose fixed for ICC. An antibody against puromycin was used for western blotting (Kerafast: EQ0001; SUnSET-WB) or immunocytochemistry (Milipore: MABE343; SUnSET-ICC) to visualise puromycin labelled proteins. Primary neurons were pre-treated in artificial cerebrospinal fluid (aCSF) for 1 hour followed by a 2 hour estradiol (10 nM) treatment. If inhibitors were relevant for the experiment, primary neurons were pre-treated for only 30 minutes and inhibitors added for further 30 minutes followed by a 2 hour estradiol treatment. Puromycin was added in the last 10 minutes of the experiment at 10 µg/mL followed by lysing for western blotting or fixing for ICC.

### Fluorescent noncanonical amino acid tagging (FUNCAT)

Primary cortical and hippocampal neurons were treated with 4 mM azidohomoalanine (AHA) AHA (Life Technologies: C10102) for 2 hours. The concentration and treatment times were verified to determine if AHA staining could be visualised within secondary and tertiary dendrites (**Supplementary Figure 1**). Primary neurons were initially pre-incubated in aCSF for 60 minutes then treated simultaneously with estradiol and AHA; these were incubated for 2 hours at 37°C with 5% CO_2_. Primary neurons were thereafter briefly washed in PBS-MC (PBS pH 7.4, 1 mM MgCl_2_; 0.1 mM CaCl_2_) followed by a 10 minute PFA-sucrose fixation at RT. These were washed 3 x in PBS pH 7.4 then simultaneously permeabilised and blocked in PBS containing 0.1% Triton- x100 and 2% NGS for 1.5 hours at RT. The neurons were washed 3 x in PBS pH 7.8 and incubated upside down in the FUNCAT reaction mix (PBS pH 7.8, 0.2 mM TBTA, 0.5 mM TCEP, 0.2 µM Alexa Fluor 555-Alkyne tag, 0.2 mM CuSO_4_) overnight at RT. The following day, the neurons were washed face up 3 x for 10 minutes in FUNCAT wash buffer (PBS pH 7.8, 0.5 mM EDTA, 1% Tween-20) and 2 x for 10 minutes in PBS pH 7.4. The neurons were then incubated in primary antibodies overnight at 4°C and fluorescent secondaries as per the protocol described previously.

#### Structural Illumination Microscopy (SIM)

SIM imaging was performed on a Nikon iSIM super-resolution microscope at the Wohl Cellular Imaging Centre. Images were acquired using a 100x oil-immersion objective (N.A. 1.49TIRF) with 51 z-steps (z-step= 0.12 µm). Raw images were acquired and deconvolved using a 3D blind algorithm specific to the iSIM to increase resolution using the NIS-Elements Advanced Research software (Nikon, version 5.01.00). All images were exported to Fiji where maximum projections were generated and subsequently analysed for puncta size, area and co-localisation, detailed below. The image acquisition parameters were the same for all images within an experiment.

#### Image analysis

The analysis of AHA along dendrites and within dendritic spines was carried out using Metamorph. Briefly, dendrite and spine regions were identified based on GFP expression, and placed into the AHA channel and used to measure average intensity of AHA.

For the analysis of co-localisaed puncta, images stained for neuronal marker MAP2, puromycin (SUnSET-ICC) and ribosomal proteins S10 (RPS10) acquired from SIM and processed through deconvolution were measured for co-localisation in Fiji. For each cell, 50 µm in length was traced along two or more separate pieces of dendrites within the MAP2 channel. As dendritic spines cannot be visualised with MAP2 staining, 2.5 µm either side of the dendrite was traced to create ‘crude spine regions’ that would encompass all the dendritic spines along that portion of that dendrite. Specifically 2.5 µm either side of the dendrite was selected, as dendritic spine lengths have typically been reported to average between 1-1.5 µm in length in 2-3 week old cultures (Boyer *et al*., 1998; Srivastava *et al*., 2011). Capturing 2.5 µm either side of the dendrite allows for any variability within dendritic spine lengths ensuring some spines are not cut off within the analysis. Thus, the ‘dendrite region’ and the ‘crude spine region’ were all assessed for puncta co-localisation. Images in the RPS10 staining channels were thresholded and particle analysis was performed on the regions of interest (ROI) within the puromycin channel to determine how many puromycin puncta would co-localise with RPS10 puncta. Expression within a region of interest was determined by whether staining intensity was greater than 25% of the maximum staining intensity.

#### Statistics

All statistical analysis was performed on GraphPad Prism 6. For all graphs, bars represent the mean average and error bars are presented as standard error of the mean (SEM). To identify differences between vehicle and pharmacological treatments, unpaired Student’s t-test with Welch’s correction was performed; the Mann-Whitney test was chosen for non-parametric data. A one-way ANOVA was employed for comparisons between multiple conditions; Bonferroni’s post-hoc analyses were performed to correct for multiple comparisons of parametric data. The Kruskal Wallis test by ranks corrected by Dunn’s multiple comparisons test was performed on non-parametric data. An online Grubb’s test was used to identify any significant outliers (https://www.graphpad.com/quickcalcs/grubbs1/).

## Acknowledgements

This work was supported by grants from UK Medical Research Council, Grant No. MR/L021064/1 to DPS; UK Medical Research Council Centre for Neurodevelopmental Disorders (Grant No. MR/N026063/1); Royal Society UK (Grant RG130856) to DPS; Independent Researcher Award from the Brain and Behavior Foundation (formally National Alliance for Research on Schizophrenia and Depression (NARSAD) (Grant No. 25957), awarded to DPS; PR was funded by a BBSRC-iCASE studentship (BB/M503356/1); IAW was funded by an ARUK studentship (ARUK-PhD2016-4); KS was supported by a McGregor Fellowship from the Psychiatric Research Trust (Grant McGregor 97) awarded to DPS; LS is supported by the UK Medical Research Council (MR/N013700/1) and King’s College London member of the MRC Doctoral Training Partnership in Biomedical Sciences; RRRD received funds from the Coordenação de Aperfeiçoamento de Pessoal de Nível Superior (CAPES, BEX1279/13-0). We thank the Wohl Cellular Imaging Centre for their help with imaging.

## DATA AVAILABILITY STATEMENT

Primary data material can be accessed by contacting the corresponding authors.

## Author Contributions

PR, HA, RRRD, IAW, KJS, KMCP, LS, MVY and DPS performed all experiments and subsequent analysis. TEP, MVY, SJM, NJB and DPS provided resources. PR, JM, SJM, NJB and DPS designed experiments. SJM, NJB and DPS oversaw the project. PR and DPS wrote the manuscript.

## References

Akama, K.T. & McEwen, B.S. (2003) Estrogen stimulates postsynaptic density-95 rapid protein synthesis via the Akt/protein kinase B pathway. J Neurosci, 23, 2333–2339.

Alvarez-Castelao, B. & Schuman, E.M. (2015) The Regulation of Synaptic Protein Turnover. J Biol Chem, 290, 28623–28630.

Avila, J.A., Alliger, A.A., Carvajal, B., Zanca, R.M., Serrano, P.A. & Luine, V.N. (2017) Estradiol rapidly increases GluA2-mushroom spines and decreases GluA2-filopodia spines in hippocampus CA1. Hippocampus, 27, 1224–1229.

Beamish, S.B. & Frick, K.M. (2021) A Putative Role for Ubiquitin-Proteasome Signaling in Estrogenic Memory Regulation. Front Behav Neurosci, 15, 807215.

Beamish, S.B., Gross, K.S., Anderson, M.M., Helmstetter, F.J. & Frick, K.M. (2022) Sex differences in training-induced activity of the ubiquitin proteasome system in the dorsal hippocampus and medial prefrontal cortex of male and female mice. Learn Mem, 29, 302–311.

Bekinschtein, P., Katche, C., Slipczuk, L.N., Igaz, L.M., Cammarota, M., Izquierdo, I. & Medina, J.H. (2007) mTOR signaling in the hippocampus is necessary for memory formation. Neurobiol Learn Mem, 87, 303–307.

Bourke, A.M., Schwarz, A. & Schuman, E.M. (2023) De-centralizing the Central Dogma: mRNA translation in space and time. Mol Cell, 83, 452–468.

Boyer, C., Schikorski, T. & Stevens, C.F. (1998) Comparison of hippocampal dendritic spines in culture and in brain. J Neurosci, 18, 5294–5300.

Briz, V. & Baudry, M. (2014) Estrogen Regulates Protein Synthesis and Actin Polymerization in Hippocampal Neurons through Different Molecular Mechanisms. Front Endocrinol (Lausanne), 5, 22.

Costa-Mattioli, M., Sossin, W.S., Klann, E. & Sonenberg, N. (2009) Translational control of long-lasting synaptic plasticity and memory. Neuron, 61, 10–26.

David, O., Barrera, I., Gould, N., Gal-Ben-Ari, S. & Rosenblum, K. (2020) D1 Dopamine Receptor Activation Induces Neuronal eEF2 Pathway-Dependent Protein Synthesis. Front Mol Neurosci, 13, 67.

Dieterich, D.C., Hodas, J.J., Gouzer, G., Shadrin, I.Y., Ngo, J.T., Triller, A., Tirrell, D.A. & Schuman, E.M. (2010) In situ visualization and dynamics of newly synthesized proteins in rat hippocampal neurons. Nat Neurosci, 13, 897–905.

Erli, F., Palmos, A.B., Raval, P., Mukherjee, J., Sellers, K.J., Gatford, N.J.F., Moss, S.J., Brandon, N.J., Penzes, P. & Srivastava, D.P. (2020) Estradiol reverses excitatory synapse loss in a cellular model of neuropsychiatric disorders. Transl Psychiatry, 10, 16.

Fortress, A.M., Fan, L., Orr, P.T., Zhao, Z. & Frick, K.M. (2013) Estradiol-induced object recognition memory consolidation is dependent on activation of mTOR signaling in the dorsal hippocampus. Learn Mem, 20, 147–155.

Hafner, A.S., Donlin-Asp, P.G., Leitch, B., Herzog, E. & Schuman, E.M. (2019) Local protein synthesis is a ubiquitous feature of neuronal pre- and postsynaptic compartments. Science, 364.

Hoeffer, C.A. & Klann, E. (2010) mTOR signaling: at the crossroads of plasticity, memory and disease. Trends Neurosci, 33, 67–75.

Huber, K.M., Kayser, M.S. & Bear, M.F. (2000) Role for rapid dendritic protein synthesis in hippocampal mGluR-dependent long-term depression. Science, 288, 1254–1257.

Jacome, L.F., Barateli, K., Buitrago, D., Lema, F., Frankfurt, M. & Luine, V.N. (2016) Gonadal Hormones Rapidly Enhance Spatial Memory and Increase Hippocampal Spine Density in Male Rats. Endocrinology, 157, 1357–1362.

Jain, A. & Woolley, C.S. (2023) Mechanisms That Underlie Expression of Estradiol-Induced Excitatory Synaptic Potentiation in the Hippocampus Differ between Males and Females. J Neurosci, 43, 1298–1309.

Jelks, K.B., Wylie, R., Floyd, C.L., McAllister, A.K. & Wise, P. (2007) Estradiol targets synaptic proteins to induce glutamatergic synapse formation in cultured hippocampal neurons: critical role of estrogen receptor-alpha. J Neurosci, 27, 6903–6913.

Jung, Y., Seo, J.Y., Ryu, H.G., Kim, D.Y., Lee, K.H. & Kim, K.T. (2020) BDNF-induced local translation of GluA1 is regulated by HNRNP A2/B1. Sci Adv, 6.

Kramar, E.A., Babayan, A.H., Gall, C.M. & Lynch, G. (2013) Estrogen promotes learning-related plasticity by modifying the synaptic cytoskeleton. Neuroscience, 239, 3–16.

Le, A.A., Lauterborn, J.C., Jia, Y., Wang, W., Cox, C.D., Gall, C.M. & Lynch, G. (2022) Prepubescent female rodents have enhanced hippocampal LTP and learning relative to males, reversing in adulthood as inhibition increases. Nat Neurosci, 25, 180–190.

Liu, F., Day, M., Muniz, L.C., Bitran, D., Arias, R., Revilla-Sanchez, R., Grauer, S., Zhang, G., Kelley, C., Pulito, V., Sung, A., Mervis, R.F., Navarra, R., Hirst, W.D., Reinhart, P.H., Marquis, K.L., Moss, S.J., Pangalos, M.N. & Brandon, N.J. (2008) Activation of estrogen receptor-beta regulates hippocampal synaptic plasticity and improves memory. Nat Neurosci, 11, 334–343.

Luine, V., Serrano, P. & Frankfurt, M. (2018) Rapid effects on memory consolidation and spine morphology by estradiol in female and male rodents. Horm Behav, 104, 111–118.

McCarthy, J.B. & Milner, T.A. (2003) Dendritic ribosomes suggest local protein synthesis during estrous synaptogenesis. Neuroreport, 14, 1357–1360.

Mukai, H., Tsurugizawa, T., Murakami, G., Kominami, S., Ishii, H., Ogiue-Ikeda, M., Takata, N., Tanabe, N., Furukawa, A., Hojo, Y., Ooishi, Y., Morrison, J.H., Janssen, W.G., Rose, J.A., Chambon, P., Kato, S., Izumi, S., Yamazaki, T., Kimoto, T. & Kawato, S. (2007) Rapid modulation of long-term depression and spinogenesis via synaptic estrogen receptors in hippocampal principal neurons. J Neurochem, 100, 950–967.

Mukherjee, J., Cardarelli, R.A., Cantaut-Belarif, Y., Deeb, T.Z., Srivastava, D.P., Tyagarajan, S.K., Pangalos, M.N., Triller, A., Maguire, J., Brandon, N.J. & Moss, S.J. (2017) Estradiol modulates the efficacy of synaptic inhibition by decreasing the dwell time of GABAA receptors at inhibitory synapses. Proc Natl Acad Sci U S A, 114, 11763–11768.

Oberlander, J.G. & Woolley, C.S. (2016) 17beta-Estradiol Acutely Potentiates Glutamatergic Synaptic Transmission in the Hippocampus through Distinct Mechanisms in Males and Females. J Neurosci, 36, 2677–2690.

Phan, A., Suschkov, S., Molinaro, L., Reynolds, K., Lymer, J.M., Bailey, C.D., Kow, L.M., MacLusky, N.J., Pfaff, D.W. & Choleris, E. (2015) Rapid increases in immature synapses parallel estrogen-induced hippocampal learning enhancements. Proc Natl Acad Sci U S A, 112, 16018–16023.

Potier, M., Georges, F., Brayda-Bruno, L., Ladepeche, L., Lamothe, V., Al Abed, A.S., Groc, L. & Marighetto, A. (2016) Temporal Memory and Its Enhancement by Estradiol Requires Surface Dynamics of Hippocampal CA1 N-Methyl-D-Aspartate Receptors. Biol Psychiatry, 79, 735–745.

Rangaraju, V., Tom Dieck, S. & Schuman, E.M. (2017) Local translation in neuronal compartments: how local is local? EMBO Rep, 18, 693–711.

Sarkar, S.N., Smith, L.T., Logan, S.M. & Simpkins, J.W. (2010) Estrogen-induced activation of extracellular signal-regulated kinase signaling triggers dendritic resident mRNA translation. Neuroscience, 170, 1080–1085.

Schanzenbacher, C.T., Sambandan, S., Langer, J.D. & Schuman, E.M. (2016) Nascent Proteome Remodeling following Homeostatic Scaling at Hippocampal Synapses. Neuron, 92, 358–371.

Schmidt, E.K., Clavarino, G., Ceppi, M. & Pierre, P. (2009) SUnSET, a nonradioactive method to monitor protein synthesis. Nat Methods, 6, 275–277.

Sellers, K., Raval, P. & Srivastava, D.P. (2015a) Molecular signature of rapid estrogen regulation of synaptic connectivity and cognition. Front Neuroendocrinol, 36, 72–89.

Sellers, K.J., Erli, F., Raval, P., Watson, I.A., Chen, D. & Srivastava, D.P. (2015b) Rapid modulation of synaptogenesis and spinogenesis by 17beta-estradiol in primary cortical neurons. Front Cell Neurosci, 9, 137.

Sheppard, P.A.S., Asling, H.A., Walczyk-Mooradally, A., Armstrong, S.E., Elad, V.M., Lalonde, J. & Choleris, E. (2021) Protein synthesis and actin polymerization in the rapid effects of 17beta-estradiol on short-term social memory and dendritic spine dynamics in female mice. Psychoneuroendocrinology, 128, 105232.

Sheppard, P.A.S., Chandramohan, D., Lumsden, A., Vellone, D., Denley, M.C.S., Srivastava, D.P. & Choleris, E. (2023) Social memory in female mice is rapidly modulated by 17beta-estradiol through ERK and Akt modulation of synapse formation. Proc Natl Acad Sci U S A, 120, e2300191120.

Sheppard, P.A.S., Choleris, E. & Galea, L.A.M. (2019) Structural plasticity of the hippocampus in response to estrogens in female rodents. Mol Brain, 12, 22.

Shrestha, P. & Klann, E. (2022) Spatiotemporally resolved protein synthesis as a molecular framework for memory consolidation. Trends Neurosci, 45, 297–311.

Smith, C.C. & McMahon, L.L. (2006) Estradiol-induced increase in the magnitude of long-term potentiation is prevented by blocking NR2B-containing receptors. J Neurosci, 26, 8517–8522.

Smith, C.C., Smith, L.A., Bredemann, T.M. & McMahon, L.L. (2016) 17beta estradiol recruits GluN2B-containing NMDARs and ERK during induction of long-term potentiation at temporoammonic-CA1 synapses. Hippocampus, 26, 110–117.

Smith, C.C., Vedder, L.C. & McMahon, L.L. (2009) Estradiol and the relationship between dendritic spines, NR2B containing NMDA receptors, and the magnitude of long-term potentiation at hippocampal CA3-CA1 synapses. Psychoneuroendocrinology, 34 Suppl 1, S130–142.

Smith, W.B., Starck, S.R., Roberts, R.W. & Schuman, E.M. (2005) Dopaminergic stimulation of local protein synthesis enhances surface expression of GluR1 and synaptic transmission in hippocampal neurons. Neuron, 45, 765–779.

Snyder, M.A., Cooke, B.M. & Woolley, C.S. (2011) Estradiol potentiation of NR2B- dependent EPSCs is not due to changes in NR2B protein expression or phosphorylation. Hippocampus, 21, 398–408.

Srivastava, D.P., Woolfrey, K.M., Jones, K.A., Shum, C.Y., Lash, L.L., Swanson, G.T. & Penzes, P. (2008) Rapid enhancement of two-step wiring plasticity by estrogen and NMDA receptor activity. Proc Natl Acad Sci U S A, 105, 14650–14655.

Srivastava, D.P., Woolfrey, K.M., Liu, F., Brandon, N.J. & Penzes, P. (2010) Estrogen receptor ß activity modulates synaptic signaling and structure. J Neurosci, 30, 13454–13460.

Srivastava, D.P., Woolfrey, K.M. & Penzes, P. (2011) Analysis of dendritic spine morphology in cultured CNS neurons. *J Vis Exp*, e2794.

Srivastava, D.P., Woolfrey, K.M. & Penzes, P. (2013) Insights into rapid modulation of neuroplasticity by brain estrogens. Pharmacol Rev, 65, 1318–1350.

Steward, O. & Schuman, E.M. (2001) Protein synthesis at synaptic sites on dendrites. Annu Rev Neurosci, 24, 299–325.

Tanaka, J., Horiike, Y., Matsuzaki, M., Miyazaki, T., Ellis-Davies, G.C. & Kasai, H. (2008) Protein synthesis and neurotrophin-dependent structural plasticity of single dendritic spines. Science, 319, 1683–1687.

Taxier, L.R., Gross, K.S. & Frick, K.M. (2020) Oestradiol as a neuromodulator of learning and memory. Nat Rev Neurosci, 21, 535–550.

Tuscher, J.J., Luine, V., Frankfurt, M. & Frick, K.M. (2016) Estradiol-Mediated Spine Changes in the Dorsal Hippocampus and Medial Prefrontal Cortex of Ovariectomized Female Mice Depend on ERK and mTOR Activation in the Dorsal Hippocampus. J Neurosci, 36, 1483–1489.

Vedder, L.C., Smith, C.C., Flannigan, A.E. & McMahon, L.L. (2013) Estradiol-induced increase in novel object recognition requires hippocampal NR2B- containing NMDA receptors. Hippocampus, 23, 108–115.

Wong, M. & Moss, R.L. (1992) Long-term and short-term electrophysiological effects of estrogen on the synaptic properties of hippocampal CA1 neurons. J Neurosci, 12, 3217–3225.

Woolley, C.S. (2007) Acute effects of estrogen on neuronal physiology. Annu Rev Pharmacol Toxicol, 47, 657–680.

Woolley, C.S. & McEwen, B.S. (1994) Estradiol regulates hippocampal dendritic spine density via an N-methyl-D-aspartate receptor-dependent mechanism. J Neurosci, 14, 7680–7687.

