## Supplemental Material for "Estradiol regulates local synthesis of synaptic proteome via sex-specific mechanisms"

**Supplemental Information:**

**Supplemental Methods**

**Supplemental Figures**

**Supplemental Methods**

**Primary neuronal culture transfections.** Primary rat hippocampal neurons were prepared as previously described (Srivastava *et al.*, 2011). Dissociated cells were seeded onto 0.2 mg/mL Poly-D-lysine (PDL; Sigma: P0899) coated 6-well Nunc plates or 60 mm Nunc dishes for biochemistry and RNA isolation or 18 mm coverglass (VWR international: 630-2200) for immunocytochemistry. Neurons were cultured in feeding media (All from Life Technologies: neurobasal medium, 21103049; 2% B-27™, 17504044; 0.5 mM L-glutamine, 25030024; and 1% penicillin:streptomycin, pen/strep, 15070063) and half media changes performed twice a week until desired age of experimentation days in vitro (DIV) 20-21. Neurons were transfected at DIV12 with 2µg peGFP-C1 (Clontech) using Lipofectamine 2000 (LF2K) (Life Technologies: 11668027) Following transfections, neurons were returned to their old feeding media and left for 7 days before pharmacological treatments were initiated between DIV20-21.

**Lysate preparation.** For whole cell lysates, primary hippocampal neurons were collected in ice cold Triton-lysis buffer: 20 mM Tris pH 7.2; 150 mM NaCl; 1% Triton X-100; and 5 mM EDTA pH 8 in the presence of protease and phosphatase inhibitors and incubated on ice for 30 minutes. Lysates were hereafter sonicated with 8-10 pulses at 40% power and centrifuged at 15’000 rpm for 15 minutes at 4°C and supernatant collected. For acute hippocampal slices, the dry weight was initially acquired per sample and lysed 10 x v/w in ice cold lysis buffer: 50 mM NaPO4; 40 mM NaCl, 5 mM EDTA; 5 mM EGTA; 1% Triton X-100 in the presence of the same protease and phosphatase inhibitors. Lysates were then sonicated with 20 pulses at

30% power, before being eluted for 2 hours on a rotor at 4°C. Hereafter, samples were centrifuged at 12’800 rpm for 20 minutes at 4°C and supernatant collected. The total concentration of all samples was measured at 562 nm using the Pierce Bicinchoninic acid (BCA) protein assay kit (Thermo Scientific; 10678484) and approximately 5-20 µg protein was loaded for western blotting. Neuron lysates were mixed with 2 x laemmli sample buffer (Bio-Rad Laboratories; 161-0737) with 355 mM β-mercaptoethanol and denatured for 5 minutes at 95°C. Alternatively, hippocampal slice lysates were mixed with NuPage LDS 4 x sample buffer (Thermo Scientific: NP0007) with 0.1 mM dithiothreitol (DTT) and denatured for 10 minutes at 70°C.

Crude synaptosomal preparations were made from primary hippocampal neurons. Neurons were collected in 150 µL ice cold homogenisation buffer: 320 mM sucrose; 5 mM Na4P2O7; 1 mM EDTA pH 8; and 10 mM HEPES pH 7.4 in the presence of protease and phosphatase inhibitors and homogenised by passage through a 25-gauge needle 12 times. The lysate was centrifuged at 800 x g for 10 minutes at 4°C to yield the P1 fraction (nuclear and large organelles) and S1 fraction (extra-nuclear). A fraction of S1 fraction was centrifuged at 15’000 x g for 20 minutes at 4°C to yield the P2 fraction (crude synaptosomes) and S2 fraction (cytosolic). The P2 fraction was resuspended in 30 µL ice cold RIPA buffer: 10 mM Tris pH 7.2; 150 mM NaCl; 1% Triton-x100; 0.1% SDS; 1% deoxycholate; and 5 mM EDTA pH 8 in the presence of protease and phosphatase inhibitors.

**Western blotting.** Neuronal and acute slice lysates were separated on 10% in-house made acrylamide gels through electrophoresis and transferred onto polyvinylidene difluoride (PVDF) membranes. Membranes were blocked in TBS-T (with 0.1% tween) containing 5% bovine serum albumin (BSA) (Sigma: A7906) followed by an overnight incubation of specific primary antibodies in the same blocking solution at 4°C with agitation. This was followed by secondary HRP incubation in the blocking solution for 2 hours at room temperature. Membranes were then incubated in Clarity Western ECL substrate (Bio-Rad: 170-5061) for 5 minutes before protein detection using the ChemiDoc XRS+ imaging system (Bio-Rad: 170-8265) running ImageLab™ software version 5.2.1 (http://www.bio-rad.com/en-uk/product/image-lab-software) (Bio-Rad). All proteins of interest and puromycin, for SUnSET, were normalised to the housekeeper β-actin whereas, the phosphorylation of proteins was normalised to its respective total protein.

**Quantification of western blots.** All images were taken in the linear range, to allow an accurate representation of the data. Acquired images were exported for analysis as a Tagged Image File Format (TIFF) and subsequently analysed using Image Studio™ Lite software version 5.2 (LI-COR). For an image, the integrated density (ID) was measured per band or smear (for Surface Sensing of Translation; SUnSET) and a background correction calculated per band. A narrow strip on the top and bottom of the box drawn around the band was used to calculate the background correction as the median of the pixels within the band. For SUnSET, the narrow strip selection was left and right of the selection box. These ID values were normalised to a housekeeper by dividing the ID value of the target protein by the ID value of the housekeeper, per condition within a biological replicate in Excel. The data set from all biological replicates within an experiment was then further normalised in the following way. Firstly, all ID values were summed for each condition across all biological replicates. Secondly, the ID values from each condition were then divided by these summed ID values. This produced a transformed set of data where each datum was relative to the sum of all conditions across the biological replicates. This method of normalisation was selected from (Degasperi *et al.*, 2014) to account for differences in antibody staining, washing and developing produced in the western blotting technique when comparing biological replicates from the same experiment across different western blots. Finally, the mean of all the vehicle ID values was taken and used to transform all the data points around the value of 1 to easily represent the fold change in protein expression after a drug treatment, for example: (1/mean of vehicles) x each datum.

**RNA isolation.** Primary neurons were collected in 1 mL TRIzol reagent (Life Technologies: 15596026) and RNA extracted according to the manufacturer's protocol. Briefly, cells lysed in Trizol were thawed at room temperature (RT) and subsequently mixed with 100 µL 1-bromo-3-phenolpropane (BCP) for 15 minutes subsequently centrifuged at 13'000 x g for 15 minutes at 4°C. The top aqueous phase was collected, and 500 µL isopropanol was added to precipitate the DNA; these were incubated at RT for 10 minutes and then centrifuged as before. RNA pellets were washed with 75% ethanol and centrifuged for 13'000 x g for 5 minutes 3 times. Pellets were left to air dry for approximately 10-15 minutes and resuspended in nuclease free H2O was added to each sample; absorbances were measured per sample using a Nanodrop spectrophotometer ND-1000.

The RNA was then treated with TURBO DNA-*free*™ kit (Thermo Fisher:AM1907) according to the company's manual. Total RNA from each extraction was incubated with 2 units of Turbo DNAse in 1 x Turbo DNAse at 37°C for 30 minutes. DNAse Inactivation Reagent was added to the RNA and incubated at RT for 5 minutes. The mixture was then centrifuged at the max speed for 3 minutes and supernatant collected. Reverse transcription was carried out on the RNA using Superscript III™ Reverse Transcriptase (SSIII) (Thermo Fisher Scientific:18080-044) according to the manufacturer's protocol. The reaction was prepared in two steps: primarily, a mixture (A) was made containing RNA, random decamers and dNTPs, which was heated to 65°C for 5 minutes and cooled on ice for approximately 1 minute. A second mixture (B) was made containing 5 X First-Strand Buffer, DTT, RNaseOUT™ Recombinant Ribonuclease Inhibitor and SSIII. This was added to mixture A producing a 20 µg final volume reaction of: approximately 907 ng RNA; 5 µM random decamers; 500 µM dNTPs; 5 mM DTT; 40 units RNaseOUT; 200 units SSIII in 1 x First-Strand Buffer. The reaction was incubated in a GS4 Thermocycler (G Storm; Pendragon Scientific Ltd.:GT-40364) following a pre-set protocol as followed: 25°C for 25 minutes; 50°C for 60 minutes; 55°C for 30 minutes; and 70°C for 15 minutes. Finally, cDNA was diluted to approximately 6 ng/µL and stored at -20°C.

**Quantitative reverse transcription polymerase chain reaction (RT-qPCR).** Expression of *dlg4* and *grin2b* was assessed using RT-qPCR. Primers were designed using Primer3 and purchased from Integrated DNA Technologies. PCR amplification of cDNA was carried out in a total volume of 12 µL using HOT FIREPol DNA Polymerase (Solis BioDyne:01-02-01000). This reaction contained 12-15ng cDNA; HOT FIREPol DNA Polymerase 1x; and 200 nM primers, and it was carried in a QuantStudio 7 Flex and the QuantStudio Real-Time PCR Software (Thermo Fisher Scientific). The following protocol was used: 95°C for 15 minutes (hot start); 40 cycles of 95°C for 20 seconds, 60°C for 20 seconds, 72°C for 20 seconds; and a continuous melting curve analysis of the amplicons from 60-95°C, which revealed single peaks, thus confirming specificity of the oligonucleotides used. Gene expression quantification was calculated relative to a standard curve of five 1:2 dilution points (quantity), and this value was further normalised to the geometric mean of the quantity of the three housekeeping/reference genes, per condition *(ACTB, GUSB, B2M)*.

**Immunocytochemistry.** Primary hippocampal neurons were washed in cold PBS and fixed with 4% paraformaldehyde in a 4% sucrose in PBS solution for 10 minutes at RT. Neurons were then washed 2 x in PBS and permeabilised and blocked simultaneously in PBS containing 0.1% Triton-x100 and 2% Normal Goat Serum (NGS; Cell signalling: 5425S) for 1 hour at RT unless otherwise indicated. Primary antibodies were added in PBS containing 2% NGS and left overnight at 4°C. The neurons were then incubated in Alexa Fluor® secondary antibodies in PBS containing 2% NGS for 1 hour at RT; dyes were selected according to species in the following wavelengths: 405; 488; 568; or 647. This was followed by a 5 minute incubation with PBS containing 0.1% the nuclear stain 4′,6-diamidino-2-phenylindole (DAPI) (Life Technologies: D1306) unless otherwise stated. Finally, the neurons were mounted with ProLong™ Gold antifade mountant (Life Technologies: P36930).

**Epifluorescence microscopy.** Epifluorescence imaging was acquired with a Zeiss Axio Imager Z1 acquired using a 40x objective (Numerical Aperture [N.A] 0.8) for puromycin optimisation (**Figure S1**). Images were exported to Fiji (https://fiji.sc/) where two-dimensional maximum projections reconstructions were generated and appropriately adjusted for brightness and contrast or pseudocoloured to emphasise areas of high staining intensities in Fiji.

**Confocal microscopy.** Confocal imaging was acquired by a Nikon A1R inverted confocal with 60x oil-immersion objective (N.A. 1.4) at the Wohl Cellular Imaging Centre (King's College London, U.K.). All images were taken as a Z-series (z-step= 0.15 µm). Images were exported to Fiji where two-dimensional maximum projections reconstructions were generated. These were further processed using MetaMorph software (Universal Imaging Corporation). Each image was background subtracted via a statistical correction. All images were taken in the linear range, to allow an accurate representation of the data. The image acquisition parameters were the same for all images within an experiment to allow comparison.

**Quantitative analysis of dendritic spine morphology and immunofluorescence.** Dendritic spine morphology and immunofluorescence analysis was carried out using MetaMorph as previously detailed (Srivastava et al., 2011) and all subsequent analysis of the data was done in Microsoft Excel. The analysis was carried out using images taken on the A1R inverted confocal with a 60x oil-immersion objective (N.A. 1.4), with at least 3 to 4 neuronal cells per condition over 4 biological replicates. Per neuronal cell, two separate pieces of dendrites, secondary and tertiary branches, were traced manually to total to approximately 100 µm in length. Using these lengths, regions were manually traced on either side of the dendrite separating the dendritic spine regions on either side along the dendrites. The dendritic spines were then thresholded by MetaMorph by outlining the spines while preserving the morphology. Each spine was then saved as a separate region to assess morphologies per condition. This method was also used to assess the average intensity of PSD-95, GluN2B and AHA within spines following estradiol treatment. The spine regions were then projected into the protein of interest/AHA channels of the corresponding image set to assess the average intensity within spines. Additionally, from the same 100 μm regions, the dendrites were thresholded by MetaMorph and the dendritic region created. The dendritic regions were then placed in the channel of interest to measure average intensity.

### Supplemental Figures

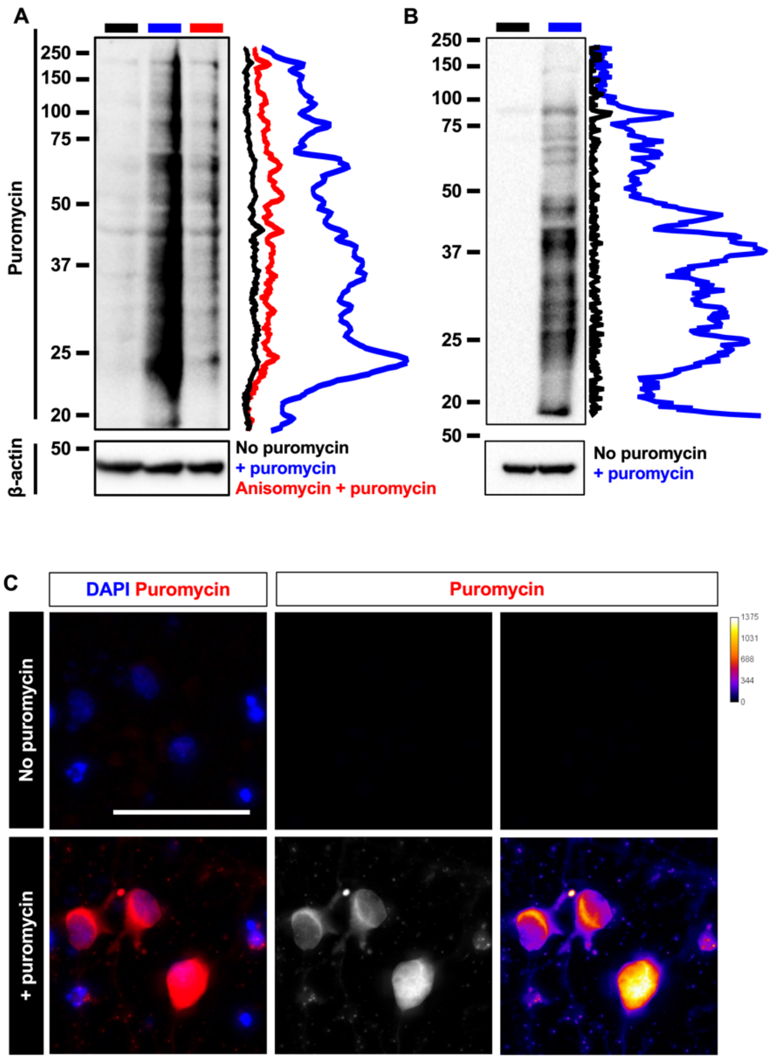


**Supplemental Figure 1.** Validation of puromycin antibodies for Surface Sensing of Translation (SUnSET) assay. **A-B** Representative western blots of days in vitro (DIV) 24 primary rat cortical neurons **(A)** and acute hippocampal slices from 10-12 week old male mouse **(B)** treated with **(A)** 10µg/mL or **(B)** 5µg/mL puromycin for 10 or 30 minutes respectively or no puromycin treatment. Primary cortical neurons were pre-treated with protein translation inhibitor, anisomycin (40µM, 30 minutes). Lysates were processed for western blotting, and immunoblotted for puromycin (Kerafast: EQ0001) and normalised to housekeeper, β-actin. In the absence of puromycin treatment, the puromycin antibody did not bind to any signal in either primary neurons **(A)** or acute slices **(B)**. This signal was reduced in the presence of protein synthesis inhibitor, anisomycin **(A)**. Line-scan next to each western blot is a visual representation of intensity changes within puromycin bands. **C** Primary cortical neurons (DIV 24) treated with 10µg/mL puromycin for 10 minutes or no puromycin treatment followed by fixation and immunostained for puromycin (Millipore: MABE343, red) and nuclei stain, DAPI (blue). No signal was observed in the absence of puromycin treatment. Scale bar = 50µm.

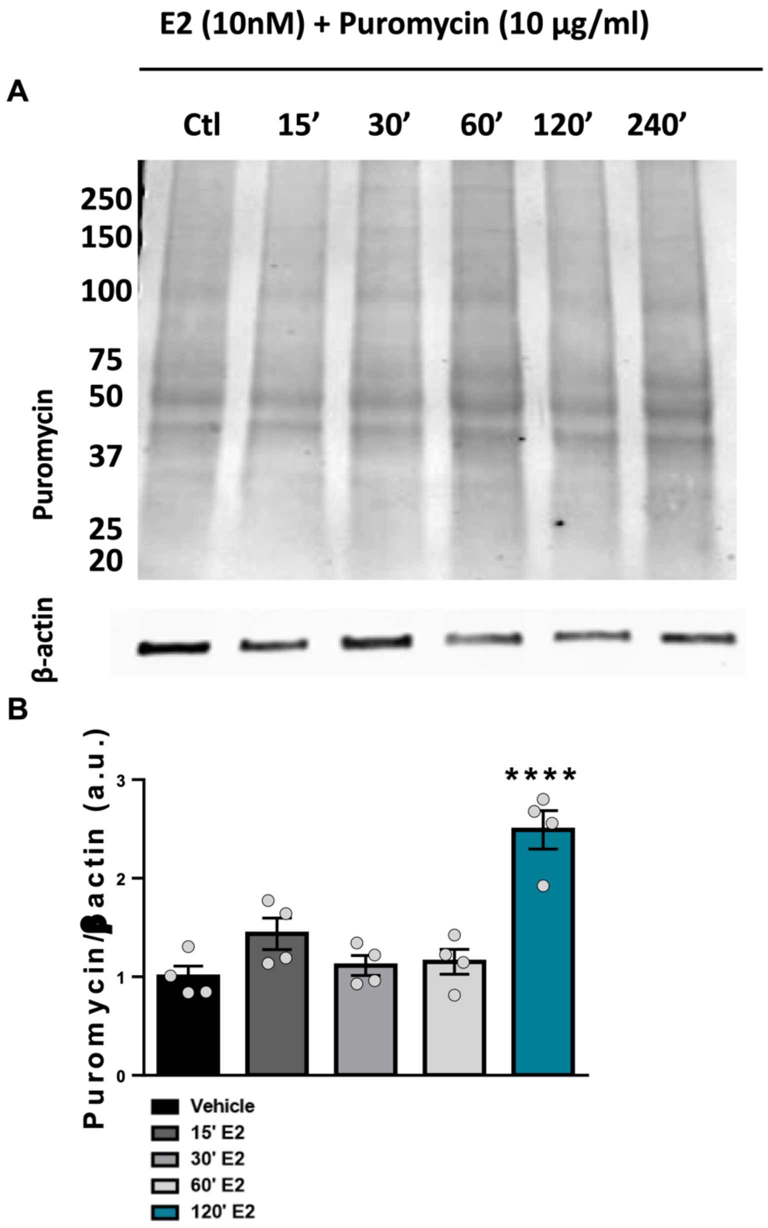


**Supplemental Figure 2.** Time-course of Surface Sensing of Translation (SUnSET) assay in response to estradiol treatment. **A** Representative western blot of days in vitro (DIV) 28 primary rat cortical neurons temporally treated with estradiol (E2; 10nM) for 15, 30, 60 and 120 minutes and 10µg/mL puromycin for 10 minutes at the end of the treatment or vehicle control (DMSO). Lysates were processed for western blotting, and immunoblotted for puromycin and normalised to housekeeper, β-actin. **B** Quantification of A. The rate of protein synthesis was increased after 120 minutes estradiol treatment. n=4 per condition over 4 different biological replicates. One-way

ANOVA, Bonferroni corrected. Error bars represent mean±SEM: **vehicle** 1.00±0.20, **15’ E2** 1.436±0.20, **30’ E2** 1.115±0.20, **60’ E2** 1.152±0.20, **120’ E2** 2.491±0.20; **** p = <0.0001.

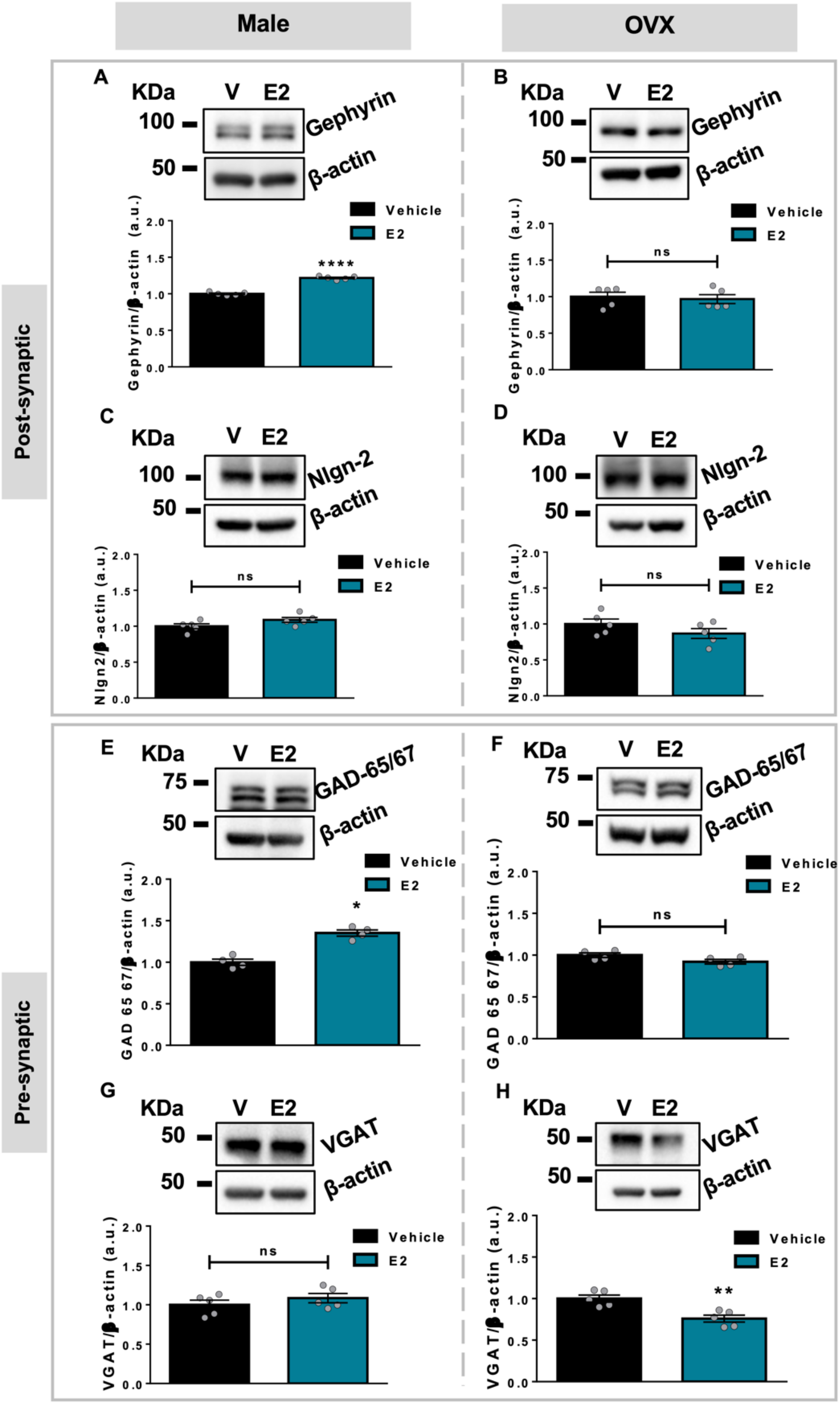


**Supplemental Figure 3.** The inhibitory synaptic proteome is regulated differentially in males and OVX females following estradiol exposure. **A-H** Representative western blots and quantification of acute hippocampal slices from 10-12 week old male **(A, C, E, G)** and ovariectomised (OVX) mice **(B, D, F, H)** treated with vehicle (DMSO; V) or estradiol (E2; 10nM) for 2 hours within the same animal. Slices were processed for western blotting, immunoblotted for post-synaptic proteins scaffolding protein gephyrin and adhesion protein neuroligin-2 (Nlgn2) and pre-synaptic proteins pre-synaptic glutamic acid decarboxylase 65 and 67 (GAD-65/67) and vesicular GABA transporter (VGAT); all were normalised to housekeeper, β-actin. Estradiol increased gephyrin expression in the male (**A,** [mean±SEM] **vehicle** 1.000±0.01, **E2** 1.218±0.01) but not OVX female (**B**, [mean±SEM] **vehicle** 1.000±0.06, **E2** 0.9673±0.06) hippocampus. No expression change was observed in Nlgn-2 levels in neither males (**C**, [mean ± SEM] **vehicle** 1.000±0.03, **E2** 1.088±0.03) nor ovx females (**D,** [mean ± SEM] **vehicle** 1.000±0.07, **E2** 0.8675±0.07) following estradiol treatment. Meanwhile, GAD-65/67 expression levels were increased in the male (**E**, [median] **vehicle** 0.9918, **E2** 1.360) but not OVX female (**F,** ([median] **vehicle** 0.9998, **E2** 0.9218) hippocampus in response to estradiol treatment. No effect was observed on VGAT expression levels in males (**G,** [mean ± SEM] **vehicle** 1.000±0.06, **E2** 1.085±0.06) but decreased expression levels in OVX females (**F**, [mean±SEM] **vehicle** 1.000±0.04, **E2** 0.7589±0.04). n=5 animals for GAD 65-67 for both male and OVX female/condition; n=5 animals for gephyrin, nlgn-2 and VGAT for both male and OVX female/condition. Unpaired student's t-tests, Welch's correction. Mann-Whitney test was used with non-parametric data sets. Error bars represent mean±SEM; * p=<0.05, ** p = <0.01, *** p = <0.001, **** p = <0.0001, ns = not significant.

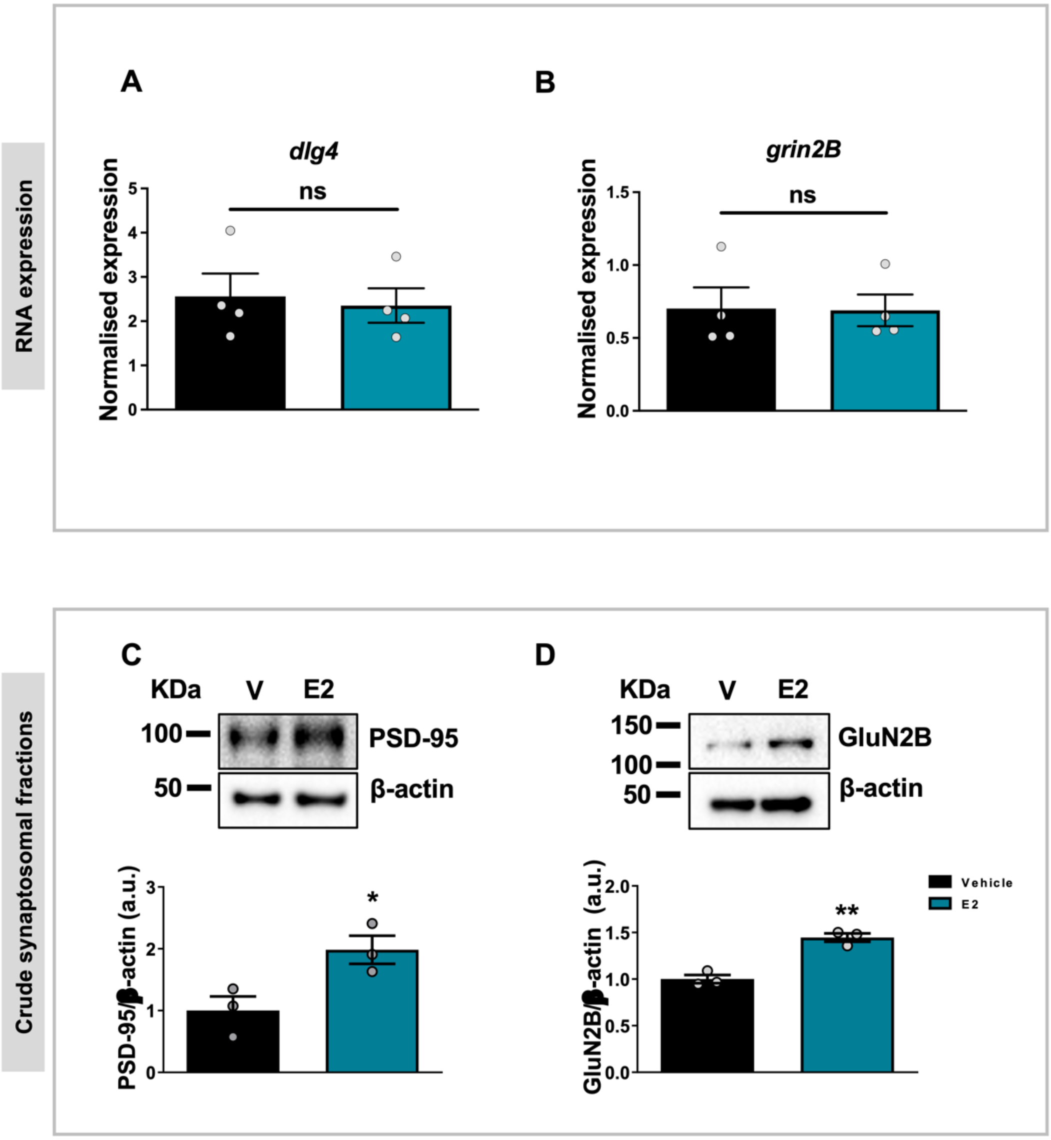


**Supplemental Figure 4.** Estradiol increases PSD-95 and GluN2B expression in crude synaptosomal fractions but has no effect on mRNA levels within the same time-frame. **A-D** Primary rat hippocampal neurons at days in vitro (DIV) 27 were treated with estradiol (E2; 10 nM) or a vehicle control (DMSO; V). **A+B** Quantification of RT-qPCR. Extracted RNA was measured for difference in DLG4 (**A,** [mean±SEM] **vehicle** 2.562±0.52, **E2** 2.353±0.394) and GRIN2B (**B**, [mean±SEM] **vehicle** 0.7017±0.15, **E2** 0.6896±0.11) expression in response to estradiol. No differences were found in the relative abundance of DLG4 and GRIN2B transcripts between estradiol or vehicle treated neurons within the same time-frame. Expression level was normalised to the geometric mean of three reference genes (ACTB, GUSB and B2M). **C-D** Representative western blots and quantification. Lysates were fractionated to separate out the crude synaptosomal fraction, processed for western blotting and immunoblotted for expression levels of PSD-95 and GluN2B and normalised to housekeeper, β-actin. Estradiol increased the expression levels of both PSD-95 (**C**, [mean±SEM] **vehicle** 1.000±0.22, **E2** 1.983±0.22) and GluN2B (**D**, [mean±SEM] **vehicle** 1.000±0.44, **E2** 1.446±0.44) within 2 hours. n=3 per condition over 3 different biological replicates. Unpaired student's t-tests, Welch's correction. Error bars represent mean±SEM; * p = <0.05, ** p = <0.001, ns = not significant.

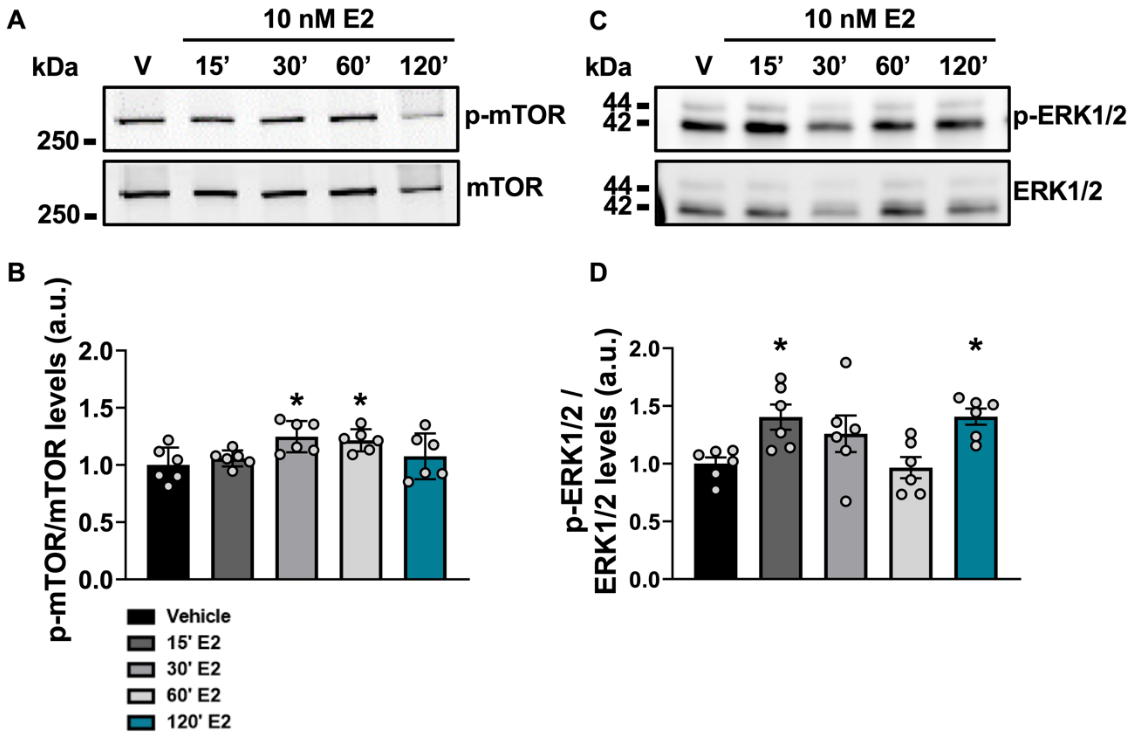


**Supplemental Figure 5.** Time-course for mTOR activation over 2 hour estradiol treatment. **A** Representative western blot of days in vitro (DIV) 24-28 primary rat cortical neurons temporally treated with estradiol (E2; 10 nM) for 15, 30, 60 and 120 minutes or a vehicle control (DMSO; V). Lysates were processed for western blotting and immunoblotted for phospho-mTOR (p-mTOR) and normalised to total-mTOR (mTOR). **B** Quantification of A. Estradiol rapidly activates mTOR within 30 miunutes, which persists until 60 minutes. The rate of protein synthesis was increased after 120 minutes estradiol treatment. n=6 per condition over 6 different biological replicates. One-way ANOVA, Bonferroni corrected. Error bars represent mean±SEM: **vehicle** 1.00±0.06, **15’ E2** 1.055±0.06, **30’ E2** 1.263±0.06, **60’ E2** 1.204±0.06, **120’ E2** 1.147±0.06; * $p = <0.05$.

Degasperi, A., Birtwistle, M.R., Volinsky, N., Rauch, J., Kolch, W. & Kholodenko, B.N. (2014) Evaluating strategies to normalise biological replicates of Western blot data. *PLoS One*, **9**, e87293.

Srivastava, D.P., Woolfrey, K.M. & Penzes, P. (2011) Analysis of dendritic spine morphology in cultured CNS neurons. *J Vis Exp*, e2794.
